# The role of promiscuous molecular recognition in the evolution of RNase-based self-incompatibility

**DOI:** 10.1101/2023.10.05.561000

**Authors:** Keren Erez, Amit Jangid, Ohad Noy Feldheim, Tamar Friedlander

## Abstract

How do biological networks evolve and expand and which parameters determine their size? We study these questions in the context of the plant collaborative-non-self recognition self-incompatibility system. Self-incompatibility evolved to avoid self-fertilization among hermaphroditic plants. It relies on specific molecular recognition between highly diverse proteins of two families: female and male determinants, such that the combination of alleles an individual possesses determines its mating partners. Though highly diverse, previous models struggled to pinpoint the evolutionary trajectories by which new alleles evolved. Here, we construct a novel theoretical framework, that crucially affords interaction promiscuity and multiple distinct partners per protein, empirical findings disregarded by previous models. We demonstrate a dynamic long-term balance between allele emergence and extinction, where their equilibrium number depends on population parameters. Our work highlights the importance of molecular recognition promiscuity to network evolvability. Promiscuity was found in additional systems suggesting that our framework could be more broadly applicable.

## Introduction

Hermaphroditic flowering plants are at high risk of self-fertilization, which would produce less fit offspring (known as “inbreeding depression” [1]). Hence, more than 100 flowering plant families have developed various mechanisms to avoid self-fertilization, generally called “self-incompatibility” (SI) [2, 3, 4, 5, 6, 7, 8, 9, 10]. Under these mechanisms, the species is sub-divided into multiple ‘types’ or ‘classes’, such that a pollen grain cannot fertilize a maternal plant of its own type. The type is encoded by a single highly polymorphic locus called the S-locus that encodes both male (Pollen-S) and female (Pistil-S) type-specifying genes.

The molecular mechanisms implementing type recognition are categorized as using either ‘self’ or ‘non-self recognition’ [6, 11, 7, 9]. Under self-recognition (SR) fertilization is by default enabled unless a maternal plant identifies an incoming pollen as having the same type as the self. Hence this mechanism requires only a single type-identifier. In contrast, under non-self-recognition (NSR), fertilization is by default disabled, and only if the incoming pollen is positively identified as having a non-self type, fertilization is unlocked. Thus, this mechanism requires multiple identifiers, that could collectively identify multiple non-self types.

Here we focus on the NSR SI mechanism demonstrated in the Solanaceae family (tomato, potato, tobacco, Petunia) and on homologous mechanisms found in Maloideae of Rosaceae (apple, pear, loquat) and Plantaginaceae (snapdragon) families, also known as the ‘RNase-based SI’. The female determinant in this mechanism is a cytotoxic S-RNase (S-locus encoded ribonuclease) [12]. S-RNase molecules expressed in female organs are imported into growing pollen tubes that attempt to fertilize the maternal plant. If the pollen is compatible, the S-RNase molecules are recognized by its male–determinant proteins and degraded, allowing for fertilization. Otherwise, if the pollen is incompatible, the S-RNases arrest the pollen tube growth, and fertilization is inhibited. The male determinant in this mechanism has been identified as an F-box protein-encoding gene and was termed S-locus F-box (SLF or SFB or SFBB) [13, 14]. Relying on non-self recognition, indeed multiple SLF genes are encoded in the S-locus to collaboratively recognize various non-self S-RNases and allow for a sufficient number of mating partners [11]. To avoid self-fertilization, a haplotype must not contain an SLF allele that recognizes its own (self) RNase, which would lead to self-compatibility. Indeed, the S-locus genes are tightly linked to avoid an accidental insertion of an SLF that could cause self-compatibility, as well as preserve the collaborative function of the entire haplotype. In the following, we refer to this mechanism as the ‘collaborative non-self recognition’ (CNSR) SI (Fig. 1).

**Figure 1:**
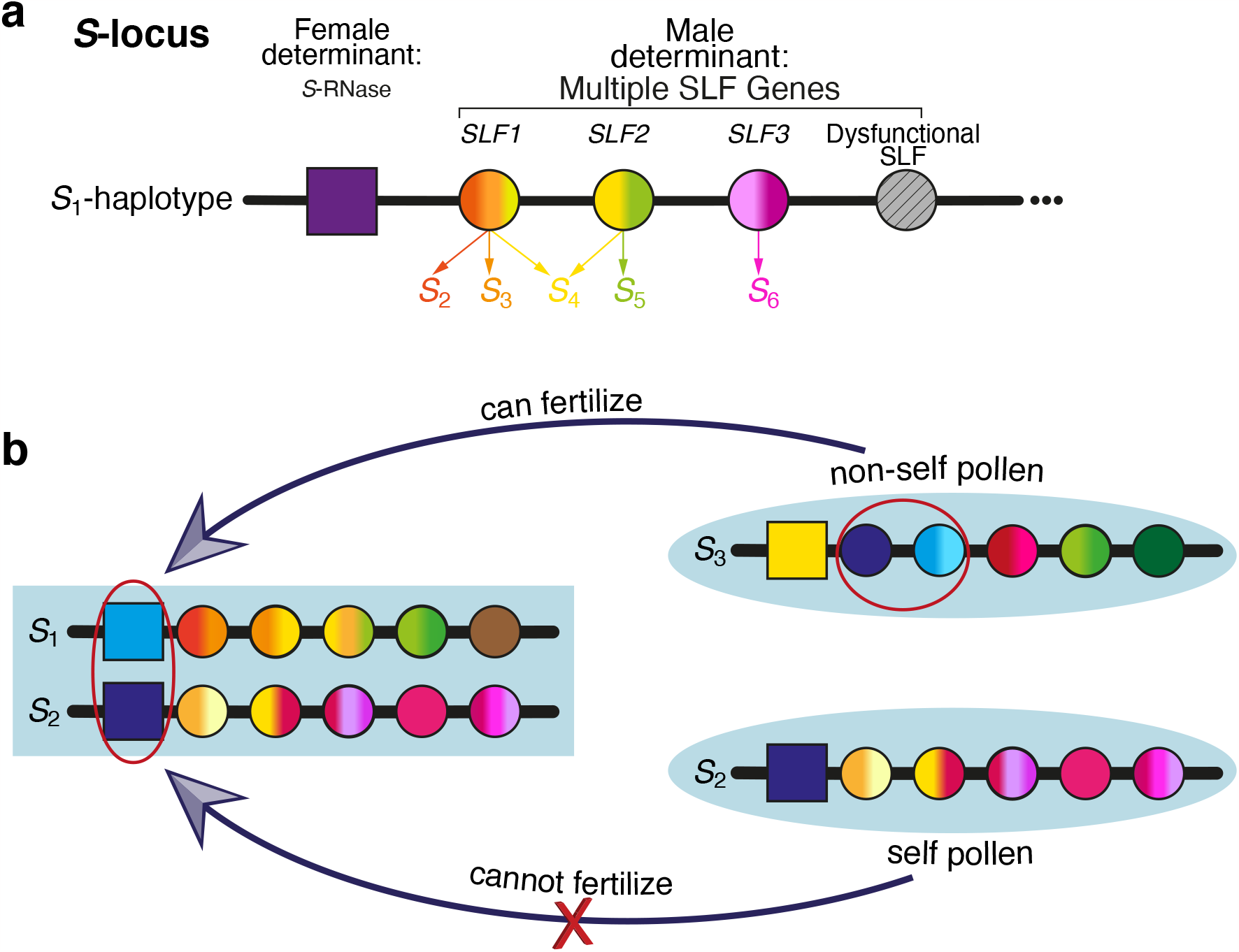
Self-incompatibility in the S-RNase based mechanism – the collaborative non-self recognition (CNSR) mechanism. **(a)** The S-locus includes a single S-RNase gene (expressed in the female organs) and multiple SLF alleles (expressed in the male organs). An SLF protein can detoxify one or more different S-RNase proteins, but could also detoxify none, in which case it is considered dysfunctional. **(b)** A haploid pollen harboring a particular combination of SLF proteins can successfully fertilize a diploid maternal plant only if it is equipped with the specific SLF alleles that can detoxify the maternal plant’s two S-RNases (top, encircled SLFs). An S-locus usually does not contain SLF alleles capable of detoxifying its own S-RNase, hence its pollen cannot fertilize its own ovules. Such a haplotype is called ‘self-incompatible’ (bottom), whereas a haplotype that does contain SLF compatible with its own S-RNase is called ‘self-compatible’. Due to inbreeding depression, there is a strong selection pressure against self-compatibility [10].

Clearly, this system requires molecular recognition that could distinguish between different molecular types, but how exactly the specificity of different RNase and SLF proteins is encoded remains elusive [15]. A combination of structural modeling [16], targeted mutations [17, 18, 19], domain swapping experiments [20, 19] and selection footprints analysis [21, 22, 23, 24] revealed that very few positions determine these protein specificities.

Despite the progress in understanding the molecular mechanisms underlying SI, a conceptual puzzle remains: while empirical evidence shows high allelic diversity, it remains murky how novel specificities evolve and which factors determine their numbers in natural populations. Early works assumed that a single mutation is sufficient to produce a new specificity [3, 25]. In these models, the equilibrium number of distinct S-alleles was determined by the balance between their introduction via new mutations and their loss by drift. Discovery of the S-locus genetic architecture led later modelers to assert that a combination of at least two mutations – in both female and male determinants – was needed to produce a new specificity. To address the conundrum of how this combination of mutations could survive, it was proposed that a new specificity could emerge via a self-compatible intermediate which later mutated to restore self-incompatibility [26, 27, 28]. Nevertheless, all the aforementioned models assumed SR-based SI, in which a newly emerging specificity was always compatible with existing ones and hence selectively advantageous owing to its rarity.

The case for CNSR is different because a new specificity is not necessarily compatible with existing ones. Thus, the process allowing new specificities to establish in the population, once they emerge, appears even more mysterious. The hurdle is that individuals carrying a novel S-RNase allele with no compatible SLF in the population are sterile as females, while novel SLF alleles that have no matching RNase yet are futile and hence their expansion in the population is only neutral. Moreover, due to the tight linkage between S-locus genes, it is insufficient that the compatible SLF appears just once, but should rather appear multiple times on all genetic backgrounds, except for the one carrying the new RNase. An earlier emergence of a compatible SLF could enable the invasion of the new RNase. Yet, in the likely occasion that this compatible SLF has only spread to part of the population, the remaining part incompatible with the new RNase is prone to extinction following the RNase invasion [29, 30]. Thus, the overall outcome of the emergence of a new RNase is not necessarily an augmentation of the total allele number but may just as well be its reduction.

These considerations place severe constraints on possible models for the evolution of new alleles in the CNSR system. Previous theoretical models that focused on the unique challenges of CNSR evolution either suggested that the novel SLF appeared multiple times on different genetic backgrounds via repeated mutations [31], or that the compatible SLF appeared just once and then spread horizontally to additional genetic backgrounds via gene conversion [7, 29]. Both models considered the scenario that the novel RNase appeared first to be deleterious and proposed that the novel SLF should precede the RNase. While these models found conditions under which new specificities could emerge, conditions for long-term allelic expansions were much more restrictive. New specificities could simply replace existing ones with no net increase in their number [31] or, even worse – cause mass extinctions of existing alleles. Hence, these models were unable to determine whether the number of distinct specificities reached an equilibrium. Previous models simplistically assumed one-to-one interactions between particular RNases and SLFs. While this is indeed the case for SR, recent empirical evidence suggested a more complicated picture for CNSR, where a particular SLF could potentially recognize several distinct RNases with possible overlaps in the recognition capacities of different SLFs [11, 32, 33, 7, 21, 34, 35].

Richer biophysical models accounting for between-residue interactions were used to study protein folding (known as lattice models) [36, 37, 38], evolution of protein-protein interaction networks [39, 40, 41], molecular recognition in receptor repertoires [42] and immune recognition [43, 44, 45]. To account for between-residue interaction energies, these models used either the Miyazawa-Jernigan potentials [46, 39, 45] or coarse-grained descriptions with only two or four amino acid categories [43, 37, 38, 40]. Similar biophysical models for protein-DNA interactions were incorporated into evolutionary models to study the evolution of gene regulation [47, 48].

Building on these evolutionary-biophysical models, we formulate a framework, which accounts for the energetic interactions between RNase and SLF proteins, and allows mating between individuals based on matches between their protein content. This framework offers several unique properties leading to surprising outcomes that were not possible before. Firstly, multiple genotypes could map into a common compatibility phenotype. Hence, a large proportion of mutations are neutral, and in particular neutral RNase mutations become possible. Secondly, our model defines a ‘compatibility-landscape’, on which genotypes that share a phenotype are connected via neutral networks and the compatibility phenotypes of neighboring genotypes are correlated, reminiscent of RNA secondary structures [49]. Thirdly, the model assumes significant promiscuity of interactions between proteins. These properties crucially allow for many-to-many interactions between RNase and SLF proteins, in agreement with empirical evidence.

Most haplotypes in our model spontaneously self-organize into ‘compatibility classes’, such that members of each class are incompatible with each other but compatible with all members of all other classes. Such classes are regularly born and die. Neutral RNase mutations not only occur regularly but turn out to be essential for the dominant class emergence trajectories. Owing to promiscuity, the model exhibits a dynamic balance between class birth and death and shows a stable equilibrium in their number. These behaviors prevail under a broad range of parameters. We propose that these are key features of the natural system, and that such evolutionary trajectories may offer a solution to the conundrum of the evolution of new SI specificities in the CNSR system. Below, we describe our model in detail, show the various trajectories for class birth and death, and demonstrate their dynamics, as found in simulations.

## Results

### An evolutionary biophysical model for the formation and extinction of compatibility classes

We consider a population of N diploid individuals, each composed of two haplotypes. Every haplotype contains a single RNase encoding the female-specificity, and multiple SLF alleles, encoding the male-specificity – Fig. 1a. Every diploid individual plays the role of a diploid maternal plant, as well as produces two types of haploid pollen carrying either of its two haplotypes. A haploid pollen can successfully fertilize a diploid maternal plant only if it is equipped with the appropriate SLF alleles that can successfully detoxify the maternal plant two RNases – Fig. 1b. It is then considered ‘compatible’ as a sire with both maternal haplotypes. Compatibility between two haplotypes could, in general, differ from their compatibility when their sire-dam roles are switched. We distinguish between unidirectional and bidirectional (in)compatibility, where the latter means the two haplotypes are (in)compatible in both roles.

For simplicity, we construct the model directly in the protein domain. We represent every allele (RNase or SLF) by a sequence of *L* amino acids, standing for the binding domain of the protein encoded by that allele. We use a size-4 alphabet representing four biochemical classes of amino acids: hydrophobic (H), neutral polar (P), positively charged (+) and negatively charged (−). We define the total interaction energy between an RNase *R*_*i*_ and an SLF *F*_*j*_ as the sum of pairwise interaction energies between their corresponding residues 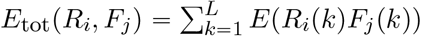 using values from [38]. These proteins are then considered matching if the total interaction energy is below a preset threshold value (Fig. 2)

**Figure 2:**
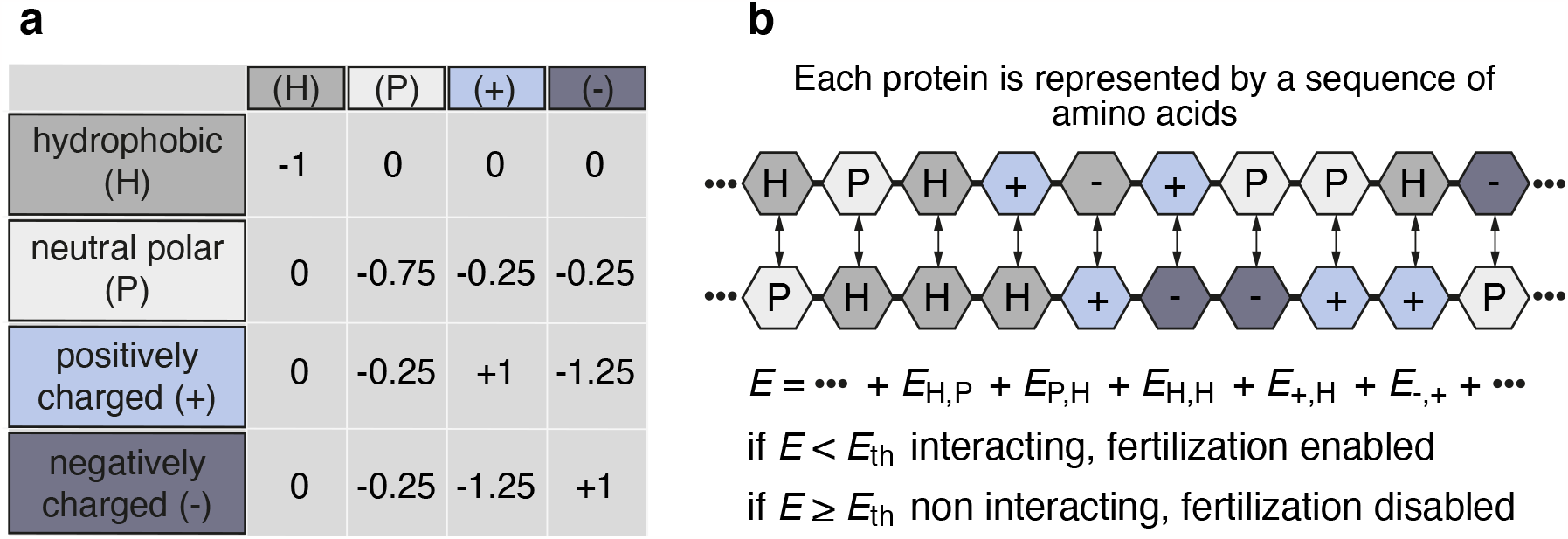
The biophysical protein-protein interaction model. **(a)** We classify the different amino acids into four categories: hydrophobic (H), neutral polar (P), positively charged (+), and negatively charged (−). We use the interaction energies between pairs of amino acids, as in [38]. **(b)** The protein-protein interaction model: we assume that the total interaction energy between two proteins is the sum of the pairwise interaction energies between their corresponding amino acids. A pair of RNase and SLF proteins are considered to be interacting (fertilization is enabled) only if this energy is below a threshold value *E*_*th*_. Otherwise, if *E* ≥ *E*_th_ they are considered non-interacting, and fertilization is disabled.

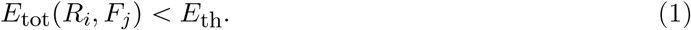

We tested several values of *E*_*th*_ (Figs. S4, S5) and *L* (not shown) and verified that the qualitative behavior of the model is persistent. For most of the results described below, these parameters were chosen following interaction energy distributions as in [50].

By construction, this model enables distinct sequences to have different numbers of matching partners, or possibly not to have such partners at all. Note also the intricate role of mutations in this model, where not every mutation essentially alters compatibility.

We simulated the evolution of a population of N such diploid individuals with the following life cycle (Fig. 3). Every generation, each of the SLF alleles is duplicated with probability *p*_dup_ and deleted with probability *p*_del_, allowing for variation in the number of SLF alleles per haplotype. Each residue in any of the RNase or SLF alleles can then be mutated with probability *p*_mut_.

**Figure 3:**
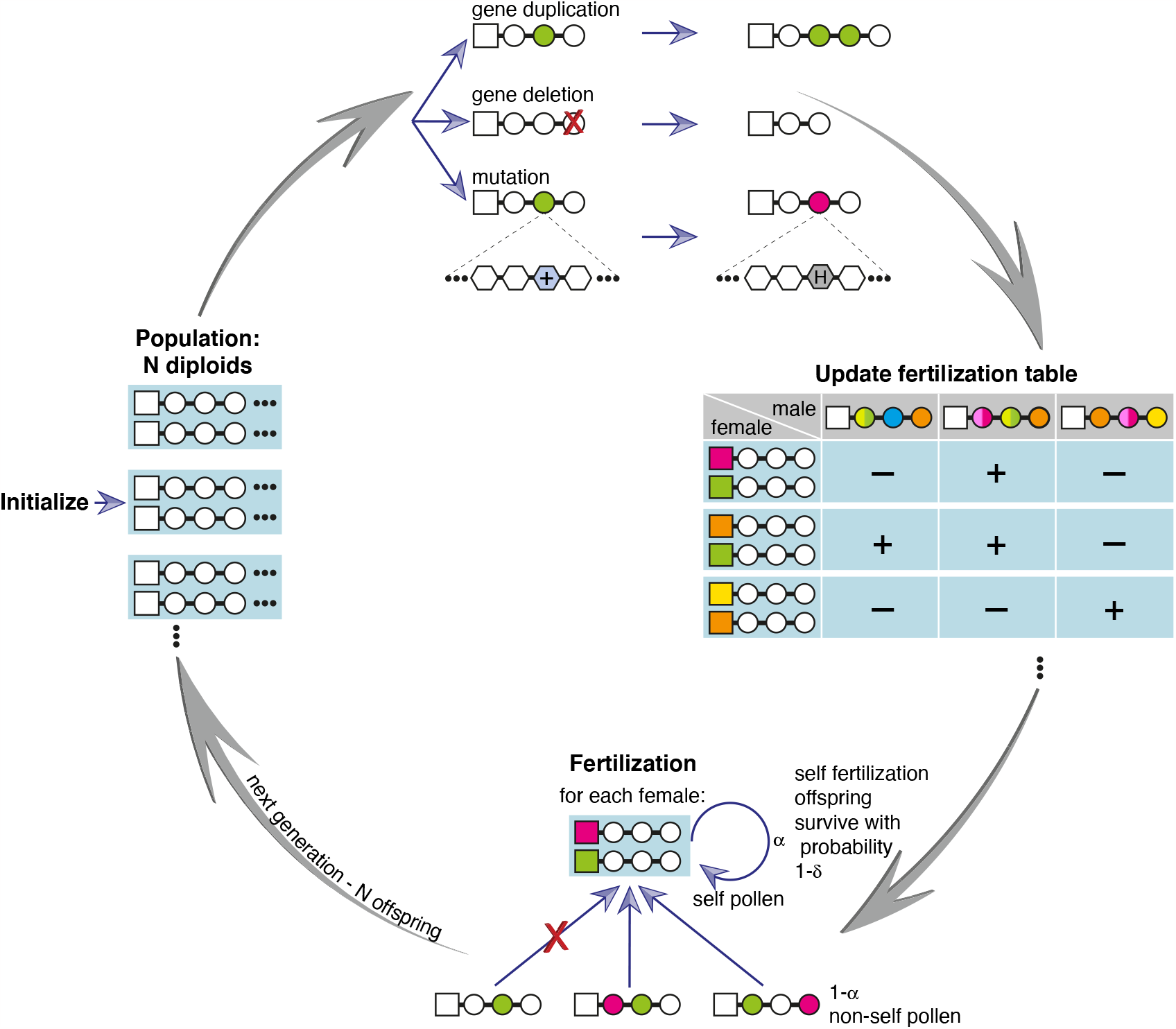
The population life cycle in our simulation. We initialize a population of N self-incompatible diploid heterozygous individuals, where each haplotype is composed of a single RNase (square) and multiple SLF alleles (circles), and every allele is represented by a sequence of *L* amino acids (Fig. 2). Every generation, each of the SLF (but not RNase) alleles could be duplicated, or deleted and each of the alleles could be mutated via substitution of randomly chosen residues. A diploid maternal plant can be fertilized by a haploid pollen, only if the pollen is equipped with suitable SLFs that can detoxify the two maternal S-RNases. To keep track of all compatible pairs, we chart the table of possible crosses between all haploid pollen and diploid maternal plant combinations. We assume that a proportion *α* of the pollen received by each maternal plant is self-pollen and the remaining 1 *α* proportion is foreign pollen. If an individual is self-compatible, only a proportion 1 *δ* of the offspring produced by self-fertilization survives. We then draw the next generation of the population by randomly picking maternal plants (with replacement) and then granting each *k* opportunities to match a randomly chosen pollen. If a matching pollen is found within *k* attempts, the maternal plant and the first successful pollen produce one offspring. This process is continued until a population of N offspring is formed, which then replaces the parental population. Each cycle represents a single generation. The default parameter values are: population size N = 500, per-residue mutation rate *p*_mut =_ 10^−4^ per generation, gene duplication and deletion rates *p*_dup=_ *p*_del_= 10^−6^ per-allele per-generation, number of generations 100,000-150,000, α= 0.95, *δ* = 1.

A haploid pollen can successfully fertilize a diploid maternal plant only if its SLFs can collaboratively detoxify both maternal plant RNases, following Eq. (1). To produce the population next generation we repeatedly pick a maternal plant uniformly at random (with replacement). If both maternal plant haplotypes are self-incompatible, we grant it multiple opportunities to mate with uniformly chosen non-self pollen grains. This sire-dam asymmetry represents the typically higher abundance of pollen grains relative to ovules. The first compatible pollen produces one offspring with one of the maternal plant two haplotypes. If any of the maternal plant haplotypes are self-compatible, it can also be self-fertilized with a certain probability, but the resulting offspring survives with a lesser probability (relative to outcrossing offspring), due to inbreeding depression (Methods).

We repeat this procedure until *N* offspring are formed. Finally, the offspring population replaces the parental population, completing one generation. See list of default parameter values in Table 1.

**Table 1:** Notation and default parameter values in the simulation.

| Parameter | Description |
| --- | --- |
| $N = 500$ | population size |
| $n_h = 10$ | initial number of haplotype types |
| $n_h - 1 = 9$ | initial number of SLF alleles in a haplotype |
| $L = 18$ | number of residues in each allele |
| $E_{th} = -6$ | interaction energy threshold below which an RNase and SLF are considered interacting |
| $p_{dup} = 10^{-6}$ [1/generation] | gene duplication rate |
| $p_{del} = 10^{-6}$ [1/generation] | gene deletion rate |
| $p_{mut} = 10^{-4}$ [1/generation] | per residue mutation rate |
| $\alpha = 0.95$ | proportion of self pollen received by a diploid maternal plant |
| $\delta = 1$ | proportion of offspring resulting from self-fertilization that do not survive |
| $k = 5$ | the number of fertilization attempts per diploid maternal plant |

### Dynamic balance between emergence and extinction of genetically heterogeneous compatibility classes

We ran the stochastic simulation multiple times for 100,000-150,000 generations each. The analyses below include only times beyond the point that the entire population descended from a single ancestral haplotype, in which its distribution is nearly independent of the initial conditions used.

We classified the vast majority of the 2*N* population haplotypes into compatibility classes, defined such that every pair of classified haplotypes were bidirectionally incompatible if they were in the same class, and bidirectionally compatible if in different classes. Importantly, classes are defined only based on the compatibility phenotype and hence class members could be genetically heterogeneous (Fig. 4).

**Figure 4:**
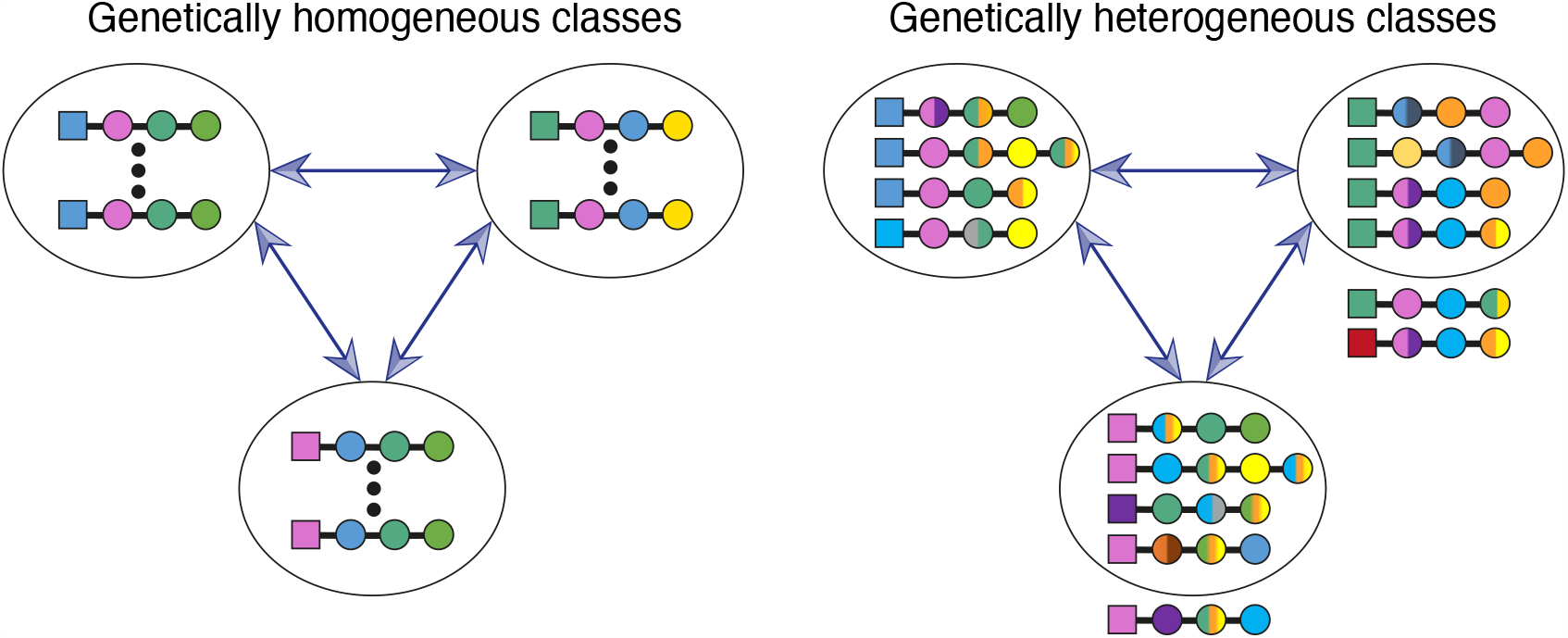
Compatibility classes are genetically homogeneous under one-to-one interactions, but could be genetically heterogeneous under the many-to-many interaction model. Compatibility classes are defined such that all members of each class are bidirectionally-incompatible within the class, but simultaneously bidirectionally-compatible with all members of all other classes. Previous models assuming that interactions between RNase and SLF are one-to-one, implied that compatibility classes should be genetically homogeneous, except for useless SLF alleles (left). Here, in contrast, we incorporate a more intricate interaction model, in which each protein could potentially have multiple different matching partners. Hence, within-class genetic heterogeneity becomes possible (right). Crucially, haplotypes in the same class could differ not only in their SLF, but also in their RNase alleles. Following this definition, some haplotypes might remain unclassified. We illustrate a few examples of unclassified haplotypes (external to the ellipses): an RNase (red) that does not have a matching SLF in any of the classes, a self-compatible haplotype (green RNase and green SLF), and a haplotype bearing an SLF that matches other class members (purple SLF).

While class definition does not necessitate that every haplotype belongs to a class, and some may be left out, we found that on average only 5% of the haplotypes remained unclassified, and only in 9% of the instances less than 90% were classified (Fig. S3). We associated each unclassified haplotype with an ‘appendix’ of the class it most recently belonged to. This small fraction of unclassified haplotypes often remained unclassified for only short time periods, typically shifting back and forth between the class and its appendix. Nearly all unclassified haplotypes either lacked one interaction with one of the foreign classes (previously termed ‘incomplete’ [31]) or had one excess interaction with some of their former class members (Methods). The number of distinct female specificities should be at least as large as the number of classes. In the following, we focus on the number of population classes, as a proxy for the number of specificities.

Next, we asked whether the number of population classes reached a stable equilibrium value with possible fluctuations around it or alternatively oscillated between high and low values. To address this question, we analyzed the population class structure every 10 generations and calculated the distribution of the number of classes amongst different population instances (Fig. 5a). We found that this distribution is unimodal and stable in time and across different runs, and hence described the process’ stationary distribution.

**Figure 5:**
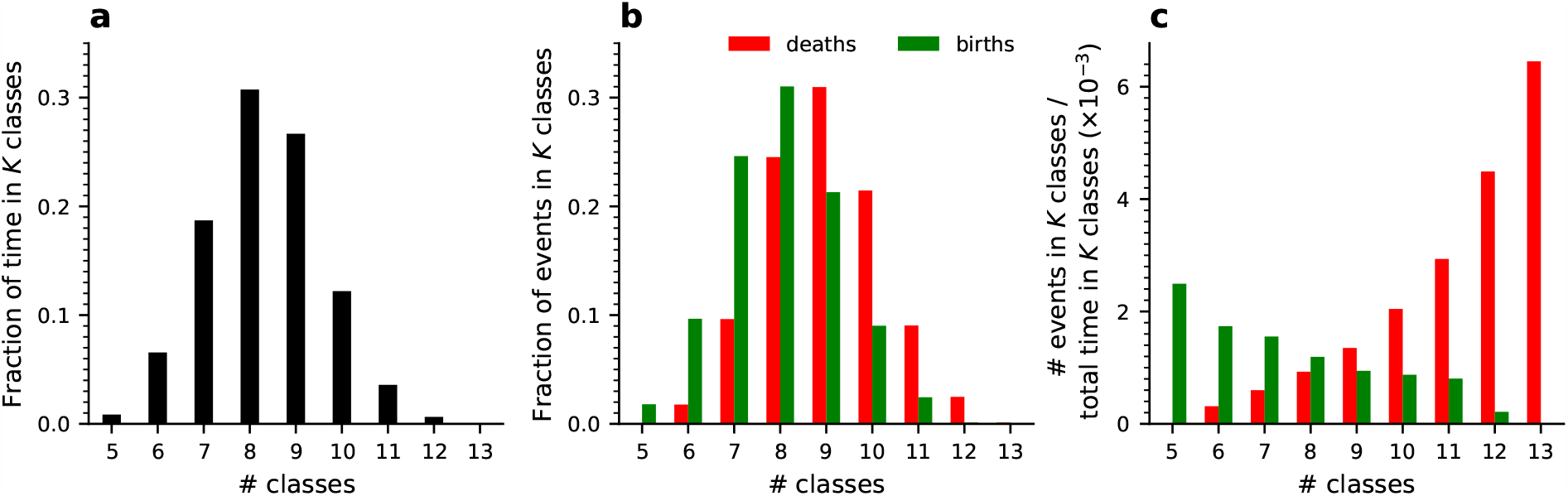
Dependence of class birth and death events on the present number of classes suggests a stable equilibrium in the number of classes – simulation results. **(a)** The fraction of time spent under each *K*-class population state. **(b)** The fraction of birth (green) and death (red) events that occurred under each *K*-class population state. **(c)** The event rates: the number of class birth (green) and class death (red) events that occurred under each *K*-class state divided by the time spent at this state. We observe opposite trends of these rates such that the class birth rate decreases but the class death rate increases with *K*, suggesting a stable equilibrium at an intermediate *K* value. The proportion of unclassified haplotypes and the proportion of self-compatible ones are shown in Fig. S3. For simulation parameters see Table 1. Results based on 39 independent runs, with a total of 1478050 generations.

To investigate the class number dynamics we traced the time points at which it either increased or decreased and designated them as class ‘birth’ and ‘death’ events, respectively (Fig. 5b). Naturally, if transitions are only possible between adjacent class numbers, the upward and downward transitions between each pair of consecutive class numbers should exactly balance each other. We observed only minor deviations from such an exact balance. Indeed, classes are always born one at a time and simultaneous extinctions of multiple classes are possible but rare.

The class birth and death rates exhibit markedly different dependence on the current class number (Fig. 5c): below the most probable class number the birth rate supersedes the death rate, and the opposite happens above this number. Hence this reversal point is a unique stable equilibrium state. This picture differs from the mass allelic extinctions, proposed in previous works [29]. We elaborate below on the reasons for this difference.

### The most prevalent split trajectories begin with a neutral RNase mutation followed by two SLF mutations

We further investigated the sequence of events leading to class birth and death. We found out that class births always occurred via the splitting of an existing class (‘mother class’) into exactly two classes. In the following we refer to class birth as ‘split’ and to class death as ‘extinction’. We identified three main routes for split and three for extinction that could sometimes intertwine, in all of which an RNase mutation played a pivotal role (Fig. 6a,b). The formation of a new class requires an RNase mutation in the mother class, followed by an SLF mutation on the background of the new RNase, which renders this haplotype compatible as sire with the original female-specificity of the mother class. To evade extinction, the latter must undergo an analogous SLF mutation rendering it compatible as sire with the novel class female-specificity. The three different trajectories described schematically in Fig. 6a correspond to the three different orderings in which the latter SLF mutation on the background of the original RNase can appear with respect to the former two mutations: after (first trajectory, blue), in-between (second trajectory, magenta), or before (third trajectory, yellow) – colors refer to the paths as illustrated in Fig. 6a. In all trajectories the first mutation is neutral, the second one confers a fitness advantage and could drive some haplotypes to extinction unless rescued by the third mutation. In principle, the three mutations needed for a split could occur in six different orderings, but the other three options pass through a self-compatible intermediate and hence are rarely seen. In Fig. 6c we show the simulation frequencies of the six optional split trajectories. These could potentially vary with the simulation parameters (see Figs. S4-S13). The two most common trajectories start with the RNase mutation as the first and neutral one. Neutral RNase mutations are a unique feature of our model, made possible by the multi-specificity of proteins. Only the third trajectory, which starts with a neutral SLF mutation, was previously studied [31], whereas the first and second ones are described here for the first time.

**Figure 6.**
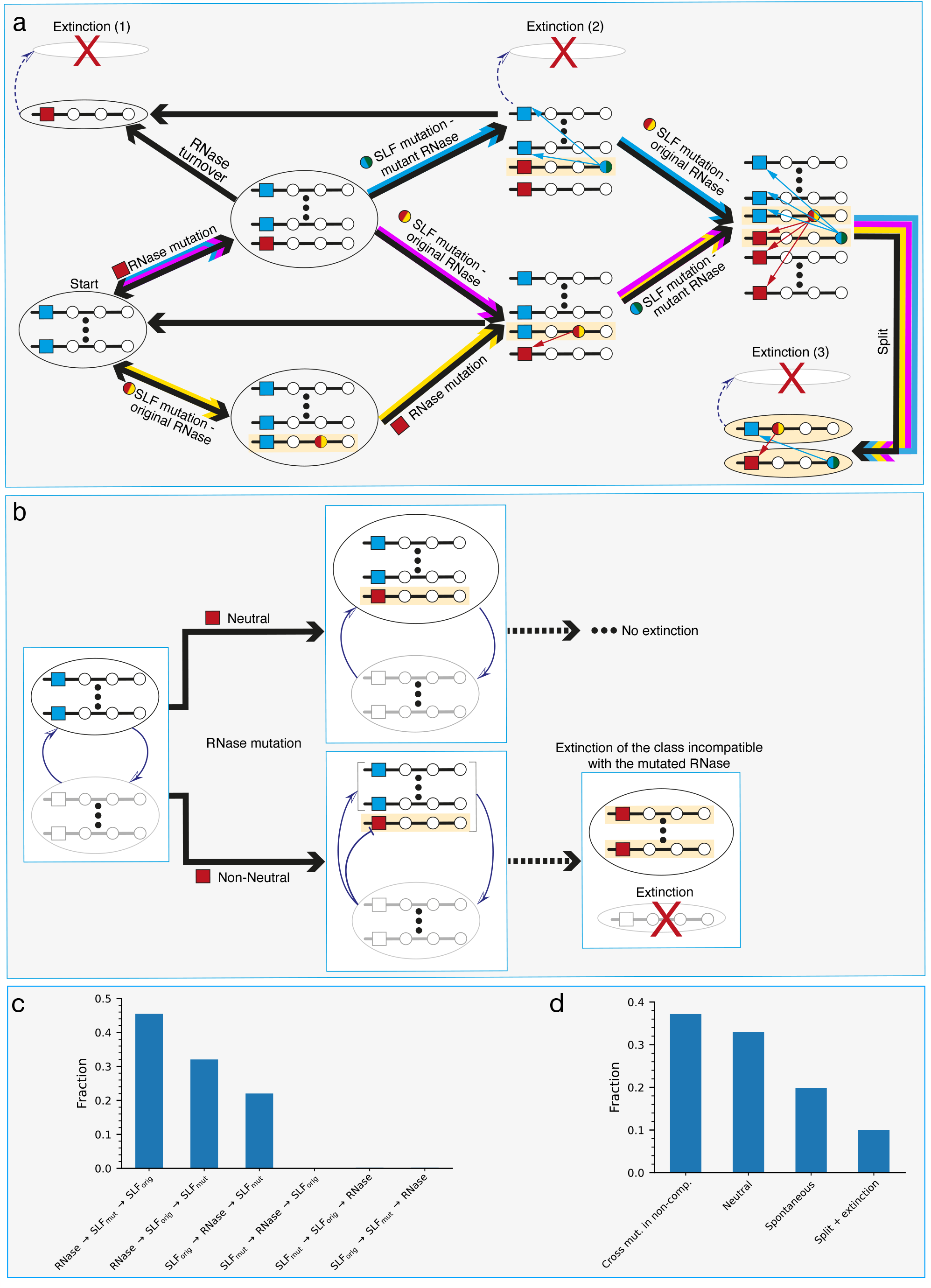
*(previous page)*: The main class split and extinction trajectories – schematic. **(a)** All split trajectories require a sequence of three mutations: one RNase and two SLF mutations. The three split trajectories – here designated by the blue, magenta, and yellow paths – differ in the mutation order. The blue trajectory starts with an RNase mutation followed by an SLF mutation on the background of the mutant RNase. Finally, a second SLF mutation compatible with the mutant RNase, which occurs on the background of the original RNase, gives rise to the daughter class separation from the mother class. The magenta trajectory is similar to the blue one, but the order of the two SLF mutations is flipped. In the yellow trajectory, the SLF mutation on the background of the original RNase precedes the RNase mutation and finally, the second SLF mutation occurs on the background of the mutant RNase. A light yellow background highlights the surviving haplotypes that form the final classes after the split. If the RNase mutation is neutral, it could only lead to a class split. If however, the newly emerging RNase mutation is incompatible as dam with one or more existing classes, it could also lead to the extinction of those classes in either of three ways, shown here as optional events (dashed arrows) accompanying the split pathways. Firstly, a neutral extinction (extinction (1)) occurs if the new RNase mutant neutrally reaches a high proportion of its class, causing fitness deficiency to the class incompatible with it as sire, till the extinction of the latter. Secondly, the spread of the mutant RNase can be accelerated by a linked SLF mutation compatible with the original RNase (extinction (2)). Thirdly, a class split could be accompanied by the extinction of another class incompatible as sire with the daughter class new RNase (extinction (3)). Split is not obligatory even if the first and second mutations occurred. For example, after the RNase mutation, it is possible that the new RNase replaces the original one and the class returns to a single-RNase state. Similarly, after the second mutation, the advantageous haplotype carrying the SLF mutation could take over before the third mutation occurs. These transitions are shown by black arrows alone. **(b)** A neutral RNase mutation (top) means that all foreign class members are compatible as sires with the new RNase (red) similar to the original RNase (blue). A non-neutral RNase mutation (bottom) means that members of at least one foreign class are incompatible with it as sires, but compatible with the original one. **(c-d)** Occurrence histograms showing the fraction of each of the possible split (c) and extinction trajectories, as observed in our simulations. In addition to the three extinction trajectories shown in (a), we also find ‘spontaneous’ extinctions, occurring with no driving mutation, solely due to drift.

In Fig. 7 we illustrate the split dynamics in three examples from our simulations, one for each trajectory type. For every trajectory, we present three plots showing (from top to bottom) the copy numbers of the relevant haplotypes *X*_*i*_, their fitness *f*_*i*_, and ‘fitness-adjusted copy numbers’ 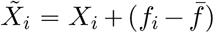. *N* (Methods) against time, where 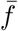 is the population mean fitness. Haplotype fitness *f*_*i*_ is defined as the proportion of diploid individuals in the population it is compatible with as sire1. The maximal fitness of a self-incompatible haplotype with *X*_*i*_ copies is 1−*X*_*i*_/*N*. If *X*_*i*_ decreases, its fitness will increase, which would later lead to an increase in its copy number, and vice versa. Hence, 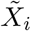 is a copy number predictor and averages out short-term fluctuations in *X*_*i*_. Specifically, negative 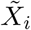 predicts the haplotype extinction. The maximal mean fitness of a *K*-class population is(*K*−2)/*K*, achieved if all classes have equal size (SI).

**Figure 7.**
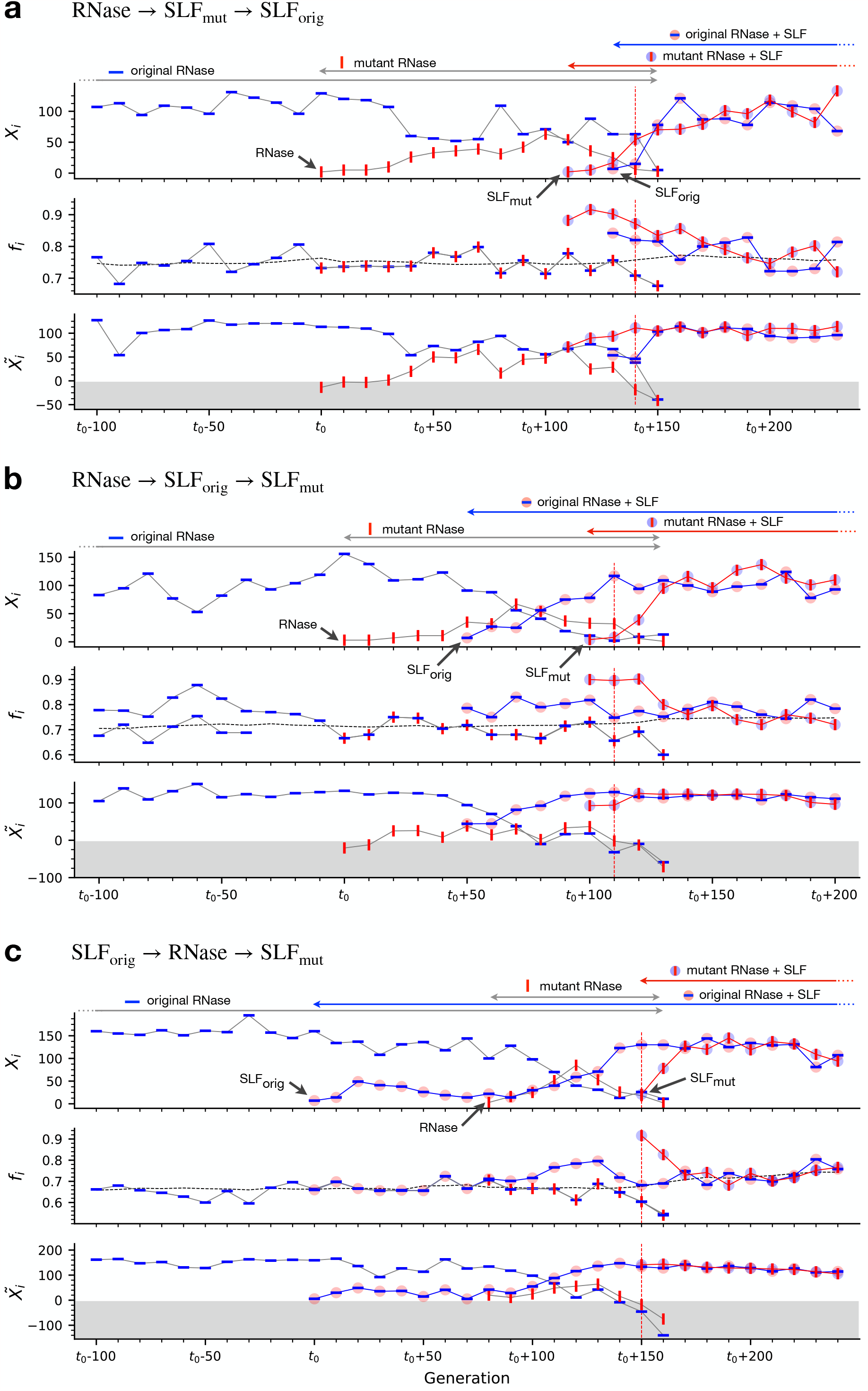
*(previous page)*: Dynamics and fitness in the three main split trajectories. Examples from simulations of the three main class split trajectories shown in Fig. 6. All examples start from a single class (‘mother-class’) and end after this class split into two classes. Each of the three panels contains three sub-figures showing: the copy numbers *X*_*i*_ of the relevant haplotypes (top), their fitness *f*_*i*_ (middle), and their fitness adjusted copy number 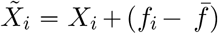. *N*, where 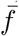 is the population mean fitness (bottom). The grey dashed lines in middle sub-figures represent the population mean fitness. Grey-colored zones in the bottom sub-figures mark negative 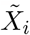 values. Haplotypes reaching negative values become extinct soon afterward. Different symbols represent different haplotypes with different combinations of mutations, as follows. The blue horizontal bar is the original RNase, red vertical bar is the mutant RNase. Colored circles surrounding the bars represent the SLF mutations, whereas the blue (red) circle represents an SLF mutation matching the original blue (mutant red) RNase. The arrows above each panel mark the lifetime of each of these haplotypes. Red dashed vertical lines in all panels show the exact split time. The trajectories differ in the order of mutations: **(a)** RNase mutation, followed by an SLF mutation on the background of the mutant RNase, and lastly another SLF mutation on the background of the original RNase. **(b)** is similar to (a) but the order of the SLF mutations is flipped. **(c)** The SLF mutation on the the original RNase background precedes the RNase mutation and lastly the second SLF mutation occurs on the background of the mutant RNase. For simulation parameters see Table 1.

Following the first trajectory, we begin with the haplotype carrying the original RNase (blue horizontal bars). Then at the time marked by t_0_ a selectively neutral RNase mutation appears (red vertical bars). As a neutral mutation, the fitness of its carriers is equal to that of the original RNase haplotypes (note overlapping curves in the fitness plot). Its initial copy number is low, but it gradually grows neutrally. If it weren’t for the SLF mutations, these two RNase variants could co-exist, until one of them would fix via drift. However, before such fixation happened, an SLF mutation, compatible as sire with the original RNase, occurred in one of the copies of the mutant RNase haplotype (red bar on a light blue circle). As the haplotype with this new SLF mutation has more siring opportunities than the other two haplotypes in its class (original RNase and mutant RNase with no SLF mutation), it immediately gains a fitness advantage over them, and its copy number sharply increases. The grey dashed line in the fitness plot (middle panel) shows the population mean fitness. Note that while the fitness values of original-RNase and mutant-RNase-only haplotypes fluctuate around this value, the new SLF-carrier fitness clearly supersedes the population mean. As all three haplotypes share a class (+appendix), an increase in one comes at the expense of the others. The initial surge of the new SLF mutant is due to the initially large copy number of the original RNase it is compatible with as sire. Later on, the decrease in the original RNase copy number reduces the SLF mutant’s fitness advantage and moderates its expansion. Lastly, the third mutation occurs – an SLF mutation on the background of the original RNase – which grants compatibility with the new RNase. Similarly, the haplotype carrying this new mutation is advantageous compared to the original RNase-only carriers. Both the original RNase-only and mutant RNase-only carriers become extinct – note that 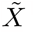 of these haplotypes go into the grey zone marking negative values. The vertical red dashed line marks the time at which the daughter class completes its separation from the mother class. The new class then grows until it equilibrates with the existing ones

The second trajectory follows similar dynamics, except that the order of the SLF mutations is reversed. Here, the SLF mutation on the original RNase background precedes the one on the mutant RNase background. As this first SLF mutant (blue horizontal bars on a red circle) has a fitness advantage over the other haplotypes not carrying it, one may think that it might lead to the extinction of the mutant RNase. However, the mutant RNase which only recently emerged is typically found in a low copy number, hence the fitness advantage conferred by compatibility with it as sire is only minor, compared to the much larger fitness advantage granted by compatibility with the original RNase, found in high copy number, in the first trajectory type. Before the mutant RNase becomes extinct, a second SLF mutation, compatible with the original RNase, emerges on its background. As before, the combination of these mutations leads to a split, while the haplotypes carrying none of the SLF mutations become extinct.

In the third trajectory, the SLF mutation on the background of the original RNase occurs first. Upon its emergence, it is neutral since the mutant RNase has not yet appeared. Once this RNase mutation appears, the already existing mutant SLF (blue horizontal bar on a red circle) is compatible with it as sire and hence its carriers gain a fitness advantage over the other haplotypes. That includes the newly emerging RNase carriers, whose copy number consequently remains at modest values. If a second SLF mutation compatible with the original RNase, occurs on one of the haplotypes carrying the mutant RNase the split will complete. The overall effect of all split trajectories is an increase in population mean fitness from 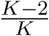 to 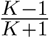 (observe the dashed line in the middle panel of each).

### Class extinctions are driven by incompatible RNase mutations

A newly emerging RNase mutation is not necessarily neutral. It might just as well be the case that it is incompatible as a dam with members of one (or more) existing non-self classes (Fig. 6b). If this happens, these classes become disadvantageous. Hence, if the new RNase expands, this could lead to the extinction of these class(es). Alternatively, since classes are genetically heterogeneous, it is also possible that if only a subset of a class is sire-incompatible with the new RNase, the class could change its composition accordingly, regain compatibility, and evade extinction. For extinction to occur, the new incompatible RNase does not necessarily need to fully replace the ancestral compatible RNase, but only to reach a sizable copy number (Fig. S14). The mutations mentioned earlier as part of class split trajectories, could just as well cause the extinction of another one, depending on the compatibility of the new RNase mutation. We identify three main extinction routes and illustrate them schematically in Fig. 6a. In Fig. 8 we show examples of the detailed dynamics and fitness of the three main extinction trajectories, in a format similar to Fig. 7. Here however haplotypes from two different classes are shown. The colored symbols are associated with the class driving the extinction and the grey circles represent the class driven to extinction.

**Figure 8.**
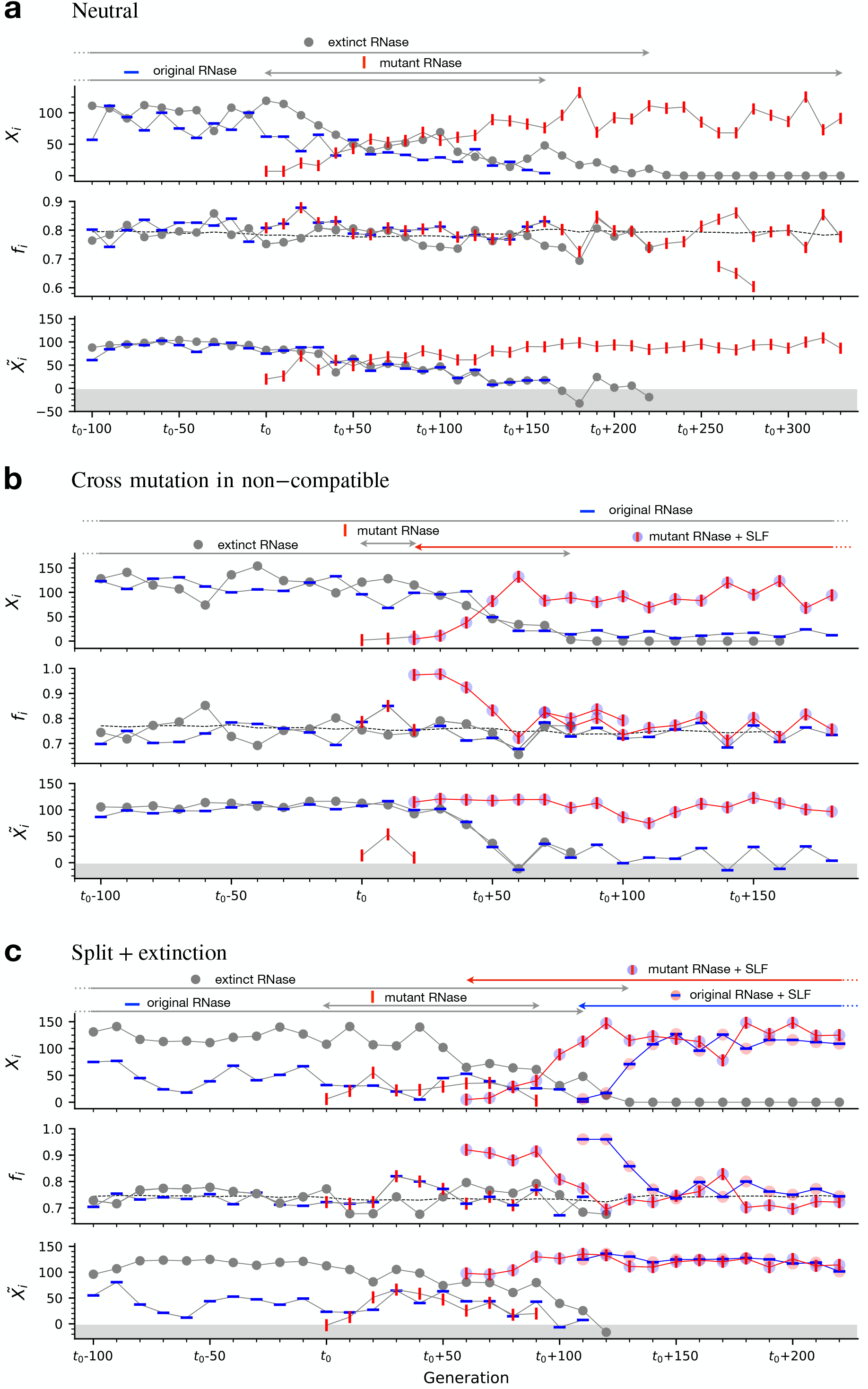
*(previous page)*: Dynamics and fitness in the three main extinction trajectories. Examples from simulations of the three class extinction trajectories shown in Fig. 6. All examples start from a state in which the class driving the extinction and the doomed class both exist and end after the doomed class becomes extinct and the driving class still exists (but potentially split). We use a similar format to Fig. 7 with 3 sub-figures for each trajectory showing the copy numbers *X*_*i*_ of the relevant haplotypes (top), their fitness *f*_*i*_ (middle) and their fitness adjusted copy number 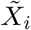 (bottom). Different symbols represent different haplotypes with different mutation combinations. The grey circles represent the doomed class. Other symbols are as in Fig. 7. The grey dashed line marks the population mean fitness. The arrows above each panel mark the lifetime of each of these haplotypes. **(a)** Neutral extinction - only an RNase mutation occurred in the driving class, which the doomed class is incompatible with as sire. **(b)** The RNase mutation in the driver class is followed by an SLF mutation on the background of the mutant RNase, accelerating its spread. This is the most common extinction trajectory. **(c)** On top of the SLF mutation of (b), an additional SLF mutation, compatible with the mutant RNase, occurs in the driving class on the background of the original RNase. This leads to a split of the driving class into two, similar to Fig. 7a. Meanwhile, the doomed class becomes extinct, because it is incompatible as sire with the newborn daughter class carrying the incompatible RNase. In the latter case the total number of classes remains intact, and hence the population mean fitness does not change. For simulation parameters see Table 1.

These extinction trajectories are the following. Firstly, the incompatible RNase mutant could expand neutrally to a sufficient number, until it causes extinction (‘neutral’, Fig. 8a). Secondly, the expansion of the incompatible RNase could be accelerated by an additional SLF mutation which confers compatibility with the original RNase (‘cross mutation in non-compatible’, Fig. 8b). Thirdly, extinction could be concomitant to a split if the daughter class RNase is incompatible with another class (‘split+extinction’, Fig. 8c). In addition, we also observe spontaneous class extinctions that are solely due to drift with no mutation involved and usually occur in very small classes (Fig. S17). In Fig. 6d we show the frequencies of the different extinction trajectories, as observed in our simulations.

To assess the likelihood of either split or extinction following an RNase mutation, we quantified the number of non-self classes that newly emerging RNase mutations (surviving more than 100 generations) are incompatible with (Fig. S18). We found that 90% of these RNase mutations are compatible with all non-self classes, hence splits are highly feasible.

While an RNase mutation could be incompatible with multiple classes, less than 1.5% of the RNase mutations are incompatible with more than one class (Fig. S18). Similarly, amongst the completed extinction events, we found that only 7% of the incompatible RNase mutations caused the extinction of two classes and only 0.6% caused the extinction of three classes, where the majority caused the extinction of only one class. Conversely, a given class could suffer from multiple incompatible RNase mutations occurring in parallel in different classes which collaboratively cause its extinction. We find that 6% of the extinctions are caused by such parallel events. To conclude, the vast majority of extinction events is driven by a single RNase mutation and lead to the extinction of one class, which differs from the picture of mass class extinctions, previously proposed [29].

## Discussion

The self-incompatibility locus is known to be highly polymorphic, but it remained obscure how allelic diversity evolved, since male and female specificity-determining genes encoded in this locus constrain each other’s evolution, posing a “chicken-or-egg” problem. While previous models proposed trajectories for the emergence of new alleles, it remained unclear whether the number of alleles reached a stable equilibrium. Here, we constructed a model for the evolution of S-alleles in the collaborative non-self-recognition SI system, which suggests a solution to this puzzle.

Our evolutionary model employs a biophysical description of interactions between the RNase (female-determinant) and SLF (male-determinant) proteins. Such models afford several desirable properties. Most importantly, they let each protein have multiple distinct partners, in agreement with empirical findings [11, 33, 51]. A biophysical description provides a natural notion of the distance between distinct proteins, thus graduating the mutation space, allowing for example, mutations that do not affect the compatibility phenotype. It further allows for promiscuous recognition between arbitrary proteins, whose extent could be tuned by varying the interaction energy threshold *E*_*th*_. Our definition of interaction between proteins induces correlation in compatibility phenotype between alleles similar in their sequences. Hence the compatibility phenotype of a mutant allele is highly correlated with that of its ancestor. This approach is in stark contrast to the more commonly used discrete-allelic models, in which only two values of distance between alleles were possible, and hence the description of mutation trajectories was over-simplified.

Our model’s attractive properties facilitate several key dynamics that were impossible in earlier models. Thanks to the inherent promiscuity of interactions, it is likely that a suite of SLF alleles is compatible with an unfamiliar RNase and even more likely if the RNase is a mutant of an ancestral RNase the SLF-suite was compatible with. Our model not only enables neutral RNase mutations, namely compatible with all existing classes, except their own, but the vast majority of RNase mutations are such. Consequently, compatibility classes can be genetically heterogeneous, not only in their SLF, but also in their RNase content. Such neutral variation amongst RNases was indeed detected [52].

A major outcome of our model is the spontaneous organization of the vast majority of population haplotypes into compatibility classes, such that there is incompatibility within class members, but full compatibility across classes. We find a total of three different paths to split and three to the extinction of classes, all of which require point mutations alone. None of these paths pass through a self-compatible intermediate, nor rely on gene conversion, as previously proposed [26, 27, 7, 31, 29, 30]. Our two most prevalent split trajectories begin with a neutral RNase mutation, and hence could not have been found in previous models, which disregarded such mutations. Only our third split trajectory begins with a neutral male-specificity mutation, similar to the fifth trajectory suggested by Bodova *et al*. [31]. However, while in [31] the neutral SLF mutation should have occurred multiple times, one at each genetic background, in our model it needs to appear just once, thanks to promiscuity.

All split trajectories require a sequence of three mutations, where the first (RNase or SLF) is neutral. The second mutation confers a fitness advantage to a sub-class (’driver’) allowing it to cross another sub-class (’endangered’). The endangered sub-class could vanish unless it gains a rescue mutation allowing it to cross back the driver sub-class and restore its fitness. We find that the time window for the rescue mutation allows for sufficient opportunities for split completion in a wide parameter range. This is partly because the fitness advantage of the driver sub-class is not constant but rather depends on the numbers of the endangered sub-class it is driving to extinction. Hence, this fitness difference is maximal upon mutation occurrence but later vanishes as the endangered population declines and the advantageous expands. In the absence of the third (‘rescue’) mutation we distinguish two phases in the sub-class dynamics following the second mutation. In the first phase, the driver is rare, hence its fitness advantage is significant and its numbers are rising while the endangered sub-class is declining. In the second phase, the driver is already prevalent and the endangered is on the verge of extinction, hence the fitness difference between them is negligible. Though unstable, the second phase allows additional time to gain the rescue mutation. For more details see mathematical analysis in SI. Rescue mutations in declining populations were studied in other contexts as well [53, 54].

We find that class extinction is driven by an RNase mutation incompatible with another existing class, in agreement with previous models [31, 29]. Yet, due to the promiscuity of interactions, most RNase mutations maintain compatibility with existing classes, and those that do not are mostly incompatible with only one class, thus extinctions typically occur one at a time (Fig. S18), in contrast to the expectations of previous models [29, 30].

Classes continuously emerge and decay, but the ratio between extinctions and splits varies with the number of existing classes K, such that for low *K* splits predominate, but for high *K* extinctions do. Thus, the class number fluctuates around a stable equilibrium, whose value positively correlates with the population size, mutation rate, and *E*_*th*_. We conjecture that extinction probability increases with the number of classes because it is affected by within-class SLF-diversity (smaller classes are less diverse and hence less resilient to new RNases) and the higher number of classes that should match each new RNase. Hence, the higher the number of classes, the lower the probability that a new RNase mutation is neutral. In contrast, if classes are fewer and hence larger, they offer both more targets for a rescue mutation and a longer time for its occurrence, because the driver sub-class takes longer to spread. Both factors increase the chances for split completion before the endangered sub-class vanishes (1st and 2nd trajectories). Larger classes also exhibit higher SLF diversity, hence a new RNase mutation is more likely to already have a compatible SLF in its own class (3rd trajectory) (Figs. S15-S16). This explanation deviates from the selection-drift balance which was proposed to determine the number of specificities in SR systems [3, 55, 56, 57] implying that a class needs to reach a minimal size to survive. In contrast, we argue that the primary barrier to class number increase is not class size, but the requirement that each class should be compatible with all others, hence a population can only tolerate a limited number of classes. Extinctions due to drift, mostly of very small classes (Fig. S17), do occur in our model but are only secondary – Fig. 6d.

Our model is robust and our main results hold under a broad range of parameter values, including varying *E*_*th*_, number of SLFs per haplotype, population size, mutation rate and various levels of inbreeding depression. Specifically, we have verified that equilibrium in the number of alleles is obtained under various parameter values (Figs. S4-S13), as long as interaction between proteins is rather likely.

Linking our model to population genetic and genomic data should be very insightful and allow testing some of the model’s qualitative predictions, as well as refining the model and calibrating its parameters. These include estimation of the number of residues distinguishing between different RNase alleles [17], and assessment of between- and within-haplotype SLF diversity [21, 58]. Our model offers predictions regarding the within-class allelic diversity and its dependence on the class size, which could then be contrasted with genomic data. We find that the number of classes should increase with population size (Figs. S6-S8), in line with previous data for SR [59]. Yet, as the quantitative relation between the population size and number of alleles could vary between SR and CNSR, it would be insightful to examine this for CNSR population data as well.

Recent experiments demonstrated that RNase allele of *Petunia* could be detoxified by various SLF alleles from different species or even different families [35]. This surprising finding is congruous with the promiscuity of interactions in our model. We also predict that RNase and SLF proteins mutated at their interaction interfaces are even more likely to interact with their pre-mutation partners than with random ones. It would be very interesting to further test this experimentally. The number of SLF alleles per haplotype determined empirically for species with CNSR SI ranged between 17 to 44 [21, 34, 60, 58, 61], which should be a proxy for the number of distinct specificities. The number of compatibility classes in our model depends positively correlates with the population and *E*_*th*_ (probability of proteins to interact at random). With 250 ≤ *N* ≤ 1000 and − 8 ≤ *E*_*th*_ ≤ − 4 the equilibrium number of classes we obtained ranged between 6 to 13 (Fig. 5, Figs. S4-S8). The larger number of classes found in natural populations could be explained by their larger sizes. Additionally, we hypothesize that population spatial structure, such that each individual has access only to a limited number of mating partners in its neighborhood, could support an even greater allelic heterogeneity beyond our current findings.

Further investigation of the model properties should address the quantitative relation between the population size and mutation rate to the number of classes, their typical lifetime, and within-class diversity. It is also interesting to determine whether the current model could be further simplified while still maintaining its current behavior.

How do proteins maintain interactions with multiple distinct partners? Do they interact with all partners via a shared interface or do they have separate or partially overlapping interfaces for different partners [40]? Here we assumed a single interaction interface per protein and did not consider shifted binding, yet it is possible that different positions in the binding interface specialize in different partners. We hope that future structural studies of these proteins will shed light on this question, and future refinement of our model could include this aspect, too. Other simplifications in our model include the lack of gene conversion and the assumption of a well-mixed population with no spatial structure. Future extensions of our model should include these as well.

Which factors determine the size of biological networks is a fundamental question applicable to different biological networks. Biochemical interactions between molecules often have relatively low energies and hence are inherently promiscuous. This promiscuity is a double-edged sword, where on one hand preserving functional fidelity could limit the network size. Indeed, avoidance of non-specific protein-protein interactions was previously proposed to limit the total number of co-existing proteins in a cell [50, 39] and regulatory crosstalk was hypothesized to limit the size of gene regulatory networks [62, 63]. Yet, on the other hand, promiscuity also grants the network additional degrees of freedom rendering it more evolvable [48, 64, 65].

In summary, we present an evolutionary-biophysical model for the evolution of allelic diversity in the CNSR self-incompatibility system. Our model is the first to allow for multiple interactions per molecule, deciphers the evolutionary trajectories of allele birth and death and resolves the question of the system equilibrium. It highlights the role promiscuous molecular recognition plays in determining the network size. Similar promiscuity may be shared by additional biological networks and the framework proposed here can be used to address further questions regarding network growth and complexity.

## Methods

### Detailed description of the simulation flow

We constructed a population-level stochastic simulation, to study the evolution of self-incompatibility alleles in the collaborative non-self recognition SI system. The simulation data analysis scripts were written in Python and run on a local server. We summarize the notation and default parameter values used in Table 1.

### Data structure

We consider a population of *N* diploid individuals, composed of two haplotypes each. Every haplotype contains a series of alleles: a single RNase encoding the female-specificity, and a variable number of SLF alleles, encoding the male-specificity.

Each allele (RNase or SLF) is represented by a sequence of *L* amino acids. Each amino acid belongs to one of four biochemical categories (hydrophobic, neutral polar, positively charged, negatively charged). We draw the sequences from the prior frequencies p_prior_ “r0.5, 0.265, 0.113, 0.122s, as obtained from the UNIPROT database (https://www.uniprot.org/) – see Table 1.

Our stochastic simulation was initiated with a population of *N* diploid individuals, each comprised of two haplotypes (described below).

### The Population Initiation module

To initialize the population we first constructed a complete set of *n*_*h*_ self-incompatible haplotypes, such that each haplotype is bidirectionally compatible (both as male and as female) with all others. We constructed this initial set of *n*_*h*_ haplotypes by the following two steps:

1. **Draw RNase sequences**. Draw an entire haplotype (one RNase and *n*_*h*_ − 1 SLFs). If this haplotype is not self-compatible, namely if none of the SLFs is compatible with the RNase – keep it. Repeat this procedure until *n*_*h*_ self-incompatible haplotypes are drawn. Then, remove the SLFs and keep only the *n*_*h*_ RNase sequences of these haplotypes. SLFs will be drawn again separately. This procedure ensures symmetry between the various RNases, such that neither is more or less constrained than the others.
2. **Draw SLF sequences with one-to-one RNase-SLF match**. Draw a single SLF sequence. If this SLF sequence is compatible with exactly one RNase from the previously drawn RNase set, place this SLF on all haplotypes except for the one whose RNase it matches. Continue this procedure, until *n*_*h*_ SLF sequences were found, such that each is compatible with exactly one unique RNase.

The outcome is a set of *n*_*h*_ complete SI haplotypes, such that each haplotype contains a single unique RNase and a set of *n*_*h*_ − 1 SLFs that are compatible with the remaining *n*_*h*_ − 1 RNases. *n*_*d*_ = *n*_*h*_·(*n*_*h*_−1) / 2 distinct diploid heterozygous genotypes can be constructed from these *n*_*h*_ haplotypes (homozygous genotypes are impossible, because all haplotypes are self-incompatible). We then draw *N* diploid heterozygous genotypes with equal probabilities.

### The simulation life cycle

The next four steps repeat every generation: gene duplication, gene deletion, mutation, and construction of the next generation, as detailed below.

1. **Gene duplication**. Each SLF (but not RNase) allele can be duplicated with probability *p*_dup_ per generation. The duplication was simulated by adding an identical allele to the haplotype sequence.
2. **Gene deletion**. Each SLF (but not RNase) allele can be deleted with probability *p*_del_ per generation.
3. **Mutation**. Each amino acid in any of the alleles (RNase or SLF) can be mutated with probability *p*_mut_ per generation. The values of amino acids chosen to be mutated are redrawn from the a priori amino acid bio-class frequencies *p*_prior_ “r0.5, 0.265, 0.113, 0.122s.
4. **Construction of the next generation**. Compute 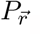, the probabilities to obtain all possible diploid genotypes 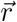 in the next generation, given their frequencies in the current generation 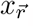, the compatibility relations between all pairs of maternal diploid genotype 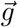 and haploid *h* and Mendelian segregation rules, namely which offspring genotype could be produced by each parental combination (using similar notation to [31]):

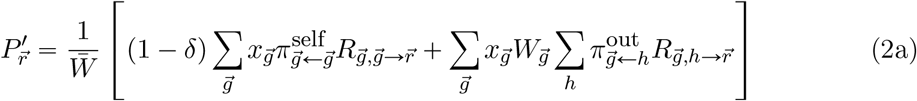

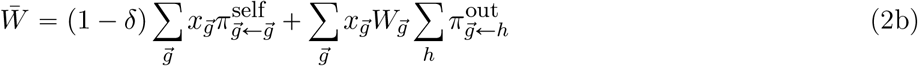

where the first term represents offspring due to self-fertilization and is non-zero only if self-compatible genotypes exist. Even if possible, these offspring survive with a lesser probability of 1 − *δ* relative to offspring resulting from outcrossing. The second term represents offspring due to outcrossing. *R* represents the probability that diploid offspring genotype 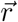 is produced by a certain parental combination 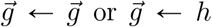 following the Mendelian segregation rules, where 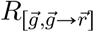 refers to self-fertilization with identical pollen and pistil genotype 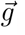, and 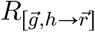 refers to outcrossing, with arbitrary diploid pistil genotype 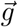 and haploid pollen genotype *h*, respectively:

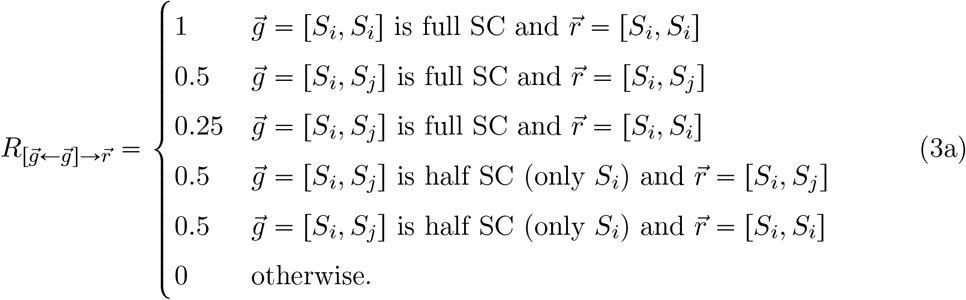

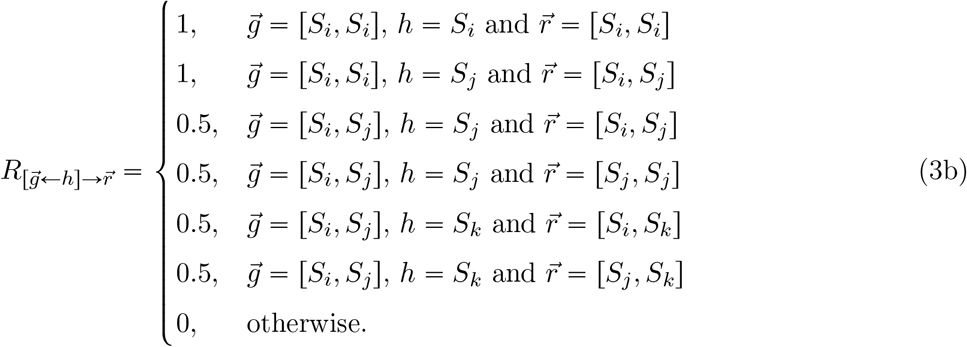

Note that outcrossing could potentially occur between genetically identical pollen and pistil, but could give rise to offspring only if they are self-compatible. 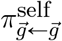 _g_and 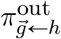 represent the probability that a maternal plant with genotype 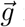 is either self-fertilized or fertilized by pollen genotype h, respectively,

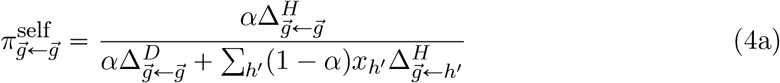

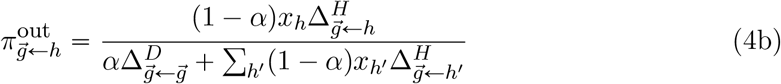

accounting for the current population frequencies of the various pollen haploid genotypes *x*_*h*_, the proportion *α* of self-pollen received, and the pollen-pistil compatibilities Δ:

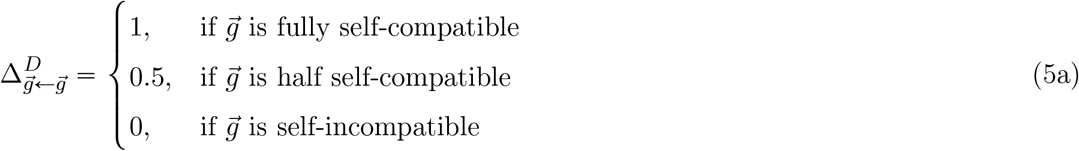

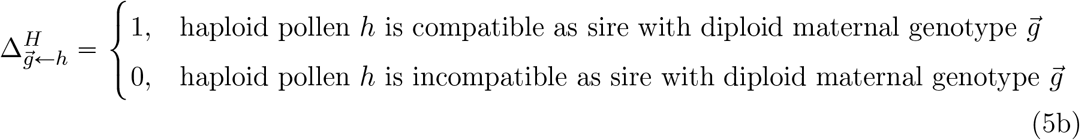

A diploid maternal plant is considered half self-compatible if exactly one of its two haplotypes is self-compatible, but the other is not. It is fully self-compatible if both haplotypes are self-compatible. Otherwise, it is self-incompatible. Once we calculated all the probabilities 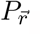, we randomly draw the genotypes of *N* diploid offspring using these probabilities. This offspring population then replaces the parental population, which completes as full generation. We then return to step (1).

To easily keep track of the possible crosses in the population we form a table of all sire-dam pairings (including self-fertilizations) and determine their compatibilities, based on their allelic contents. Every generation we update this table as necessary to account for the effect of newly emerging mutations on fertilization capabilities.

### Number of fertilization attempts

While pollen grains are typically more abundant than ovules, it is still possible that certain maternal plants suffer from pollen limitation, for example, if they possess a female specificity with too few compatible sires. To account for that we modified the fitness function to reflect a limited number *k* of fertilization attempts per maternal plant:

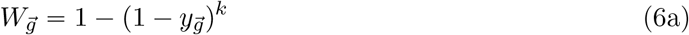

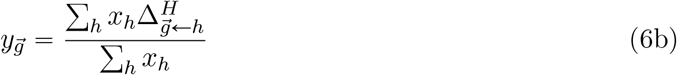

where 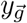 is the proportion of pollen that is compatible with diploid maternal genotype 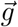 out of all foreign pollen the maternal plant receives. In practice, as our model works in a regime, where the vast majority of novel RNases are compatible with all existing pollen genotypes, this parameter had very little effect, unless extreme pollen limitation was assumed (*k* =1 to 2).

### The simulation data structure and output

We store the population state in the HAPLOTYPE data structure, which includes a list of haplotype objects. Each object in the list contains the following information: the haplotype’s current copy number, the identity of the haplotype ancestor at the initial condition, the generation in which the haplotype first emerged, the identity of its direct parent haplotype, and its allelic content, namely the list of SLFs and RNase indices it contains. This data structure only keeps the information of the current population and is updated every generation. Allelic indices are set uniquely, such that every new allele receives a unique index, not given previously to any other allele, including alleles that no longer exist in the population. If the same allele emerged independently more than once, it received the same index upon each emergence. Furthermore, the simulation keeps a list associating the allele indices with the relevant amino acid sequences for all SLFs and RNases that ever existed in the population. In addition to the current population state stored in the HAPLOTYPE structure, we also keep lists of the parental indices for all haplotypes that ever existed in the population starting from the time that the entire population descends from only original initial condition haplotypes. This information is essential when tracking the class evolution (see below) and especially detecting the split/extinction moment.

During each simulation run, we save the HAPLOTYPE data structure documenting the population state, every 10 generations for further offline analyses. We analyze the simulation data only from the time point that all the population haplotypes descend from only one of the *n*_*h*_ ancestral initial haplotypes. By that, we minimize the dependence of the results on the specific initial condition of each run. The typical run-time needed until only a single ancestor remained was ≈ 50, 000 generations. From that time-point the simulations were run for additional 100,000 generations.

### The haplotype classification algorithm

Every 10 generations, we classified the set of haplotypes in the population according to their compatibility phenotype. The classification should obey the following two requirements: a haplotype affiliated with a class should not be compatible neither as sire nor as dam with any of the other members of the same class, but should be compatible both as sire and as dam with all members of all other classes, it is not affiliated with. Following this definition, it is not guaranteed that all haplotypes are classified into any of the classes. Specifically, self-compatible haplotypes cannot be associated with any class, because they can fertilize their own class members. The division of haplotypes into classes is not unique. To ensure that the most frequent haplotypes are classified, we first ordered all the self-incompatible haplotypes in the population in descending order of their copy number. We then defined the most frequent haplotype to be the first class. Then, For every unclassified haplotype in its turn (following the descending copy number order), we tested its compatibility with the already existing classes. If it was bidirectionally incompatible with all members of exactly one class and bidirectionally compatible with all members of all the remaining classes, we associated it with the class it was incompatible with. If it was bidirectionally compatible with all members of all existing classes, we defined a new class and associated this haplotype with that class. Otherwise, this haplotype remained unclassified and we moved on to the next one. Importantly, our class definition refers only to the compatibility phenotype of haplotypes, rather than to their genotype. As our model allows for multiple partners per allele, classes could be genetically heterogeneous.

This classification procedure was repeated every 10 generations independently of previous classifications. As haplotypes mutated during the simulation, this could influence their class association. For example, if a haplotype gained an SLF mutation that rendered it compatible as a sire with (some of) its class members, it no longer followed the requirement of being incompatible with all its class members. Similarly, if an RNase mutation occurred, that was incompatible as a dam with members of foreign classes, it no longer followed the requirement for bidirectional compatibility with all non-self classes. Many of these mutations have a short lifetime, resulting in haplotypes that exit and enter the classes frequently. Such mutations are also part of the trajectory to class split and extinction. Hence, to keep the information regarding the class from which the (temporarily) unclassified haplotype originated, and the one to which it could soon return, we additionally defined class appendices.

These class appendices include former class members that following a mutation, either lost compatibility with members of foreign classes or gained compatibility with their class members or both. Hence, each haplotype is uniquely associated with either one of the classes or with one of the class appendices, but never with two of them. This definition of class appendices allows us to conveniently track class evolution.

### Identification of split and extinction events

Other possible changes in the population class structure are the split of an existing class into two separate ones and the extinction of an entire class. To identify such events we need to associate between the class structures obtained at different time points. To accomplish that, we tracked the parents in the previous classification time point of haplotypes affiliated with a certain class. If all the parents of haplotypes in that class are also affiliated with one class in the previous classification, we identify the current class with the parental class. We identify a split if the offspring of parents associated with one class in the previous classification are associated with two separate classes in the current classification. Such a split is the end of the process during which, mutated haplotypes, containing mutated RNase/ SLFs, were associated with the class appendix.

Class extinction was defined as the event that all haplotypes of a class and its appendix leave no descendants in the next classification, including no mutants of the former class haplotypes. We specifically require that the appendix members also vanish since a haplotype can alternate between being associated with a class and being associated with its appendix, depending on the state of the remaining classes.

### Haplotype grouping

In Figs. 7-8 we illustrate copy numbers and fitness of haplotypes involved in split and extinction events. There are five categories of haplotypes shown there: ‘original RNase’, ‘mutant RNase’, ‘original RNase + SLF’, ‘mutant RNase + SLF’, and ‘extinct RNase’ (see the main text for full details). Yet, each category could potentially contain multiple different haplotypes. For example, the ‘original RNase’ category refers to all haplotypes sharing that RNase, but they could vary in their SLFs content, as long as this has no implication on haplotype compatibility. Similarly, some ‘mutant RNase’ copies could later acquire SLF mutations that do not affect their functionality. For simplicity of presentation, we group together all haplotypes sharing a common functionality and show the sum of their copy numbers.

### Calculation of fitness-adjusted copy number for grouped haplotypes

For ease of presentation, we lump together distinct haplotypes with shared functionality (e.g. in Fig. 7 and Fig. 8). The fitness adjusted copy number is defined as 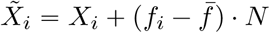, where *X*_*i*_ is the number of copies, *f*_*i*_ is the proportion of diploid maternal plants it is compatible with as sire and 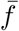 is the population mean fitness. If these different haplotypes happen to have the same fitness value, we simply add their copy numbers. For example for two haplotypes *X*_1_, *X*_2_ with equal fitness values *f*_1+ 2_ = *f*_1_ = *f*_2_:

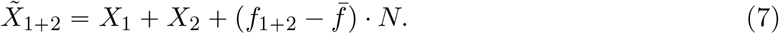

If however, they differ in their fitness, we use in the formula the weighted sum of their fitness values

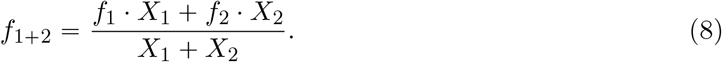

instead of *f*_*i*_.

## Acknowledgements

We thank Idan Efroni, Yonatan Friedman and Avi Mayo for comments on the manuscript and Amnon Horovitz for useful discussions. This research was supported by the Israel Science Foundation, grant number 1889/19 (T.F.).

## Footnotes

1 As we show below, most RNase mutations are compatible with most non-self classes. Hence, reproductive capacity is insensitive to the female-side fitness.

