## Supplementary Information for "The role of promiscuous molecular recognition in the evolution of RNase-based self-incompatibility"

October 5, 2023

#### The time window for the splitting process

Mutations happen continuously, but only specific combinations of three mutations could, under the circumstances described, lead to splits. In all three split trajectories, the first mutation is neutral. This mutation can possibly be lost or fixed in the population: a neutral RNase can either decay or replace the ancestral RNase and SLF mutation remains neutral unless by coincidence an RNase mutation to which it matches emerges. The second mutation in all trajectories confers a fitness advantage to the carriers of the first SLF mutation, thereby triggering its expansion. To complete the split, the third mutation should occur on a different haplotype, which is selectively inferior and under decline. Thus it must occur within a limited time window before that haplotype is lost. In Fig. S1(a-c) we show the distribution of mutation opportunities $n_\mu$ between the second and third mutations in all completed splits as observed in our simulations, dissected by trajectory type. Mutation opportunities are defined as the cumulative copy number of the sub-class waiting for the rescue mutation until it appears $n_\mu = \sum_{t=T_2}^{T_3} X(t)$, where $T_{2,3}$ are the times of the second and third mutations, respectively.

The distributions shown in Fig. S1(a-c) are based only on splits that were completed, and do not include aborted splits, due to a third mutation that did not arrive on time. To test the feasibility of split completion, we focused on the three split examples illustrated in Fig. 7 as test cases. For each, we took the population state shortly after the second mutation (to avoid the drift barrier) as the starting point. We then disabled the possibility of additional mutations and tested in repeated stochastic simulations the time till $T_\infty$ – the extinction of the selectively inferior haplotype that should receive the third mutation – conditioned on the survival of the (selectively superior) haplotype carrying the second mutation. We show here the distributions of mutation opportunities $n_\mu = \sum_{t=T_2}^{T_\infty} X(t)$, as obtained in multiple repeats all starting from the same initial population state (Fig. S1d-f). These distributions represent the feasible time for split completion.

Mutations are random independent events, hence their occurrence is often approximated by a Poisson process. Adopting this approach, we can obtain rough estimates for the typical number of mutation opportunities required for split completion using the numbers obtained in our simulations. Using Fig. S1a we estimate $\lambda$ the rate at which split-completing-mutations occur to be at least one such mutation per 3600 mutation opportunities (the rate is likely higher, because all the cases in which the mutation occurred earlier could have afforded additional mutations). Using the mean numbers of mutation opportunities obtained in Fig. S1d-f, the probabilities that no mutation occurred in that number of mutation opportunities $\langle n_\mu \rangle$ is $\exp^{-\lambda\langle n_\mu \rangle}$. Hence the probability that at least one such a mutation occurred and the split completed is $1 - \exp^{-\lambda\langle n_\mu \rangle}$. Using the numbers obtained in Fig. S1d-f, these probabilities are 0.64, 0.36 and 0.08, for the three trajectories, respectively. This estimate supports our finding in 6c that the first trajectory is the most common. Indeed, in Fig. 6c we show that the first trajectory forms 45% of the cases and the second trajectory only 33% of the cases, though the ratio is more modest than suggested by our estimate. The probability for split completion in the third trajectory is significantly lower compared to the other two, but it is partly compensated by a much larger number of split initiations, because it starts with an SLF mutation, rather than an RNase mutation, as the other two trajectories. The much larger number of SLFs compared to RNases and the lower penalty on SLF evolution render the initial event of SLF mutation much more likely. Hence, the split completion probability alone can only be used to compare the occurrences of the first and second trajectories, both starting with an RNase mutation, but not with the third. As a caveat, we also note that Fig. S1d-f show each one particular case, that might not be characteristic of the entire distributions shown in Fig. S1a-c.

### Mathematical treatment of the model

In this section, we mathematically analyze a simplified version of the full model used in the simulations. Throughout the analysis we do not allow mutations, so that new haplotypes cannot emerge in the course of the evolution. While admittedly simplified, it retains the main properties and behavior of the full model. This analysis extends and generalizes results for the haploid model obtained by Iwasa and Sasaki [1].

#### The model equilibrium points

Throughout we introduce two simplifications. Firstly, we assume that all haplotypes in the population fall into one of $\ell$ compatibility classes, such every two haplotypes are bidirectionally compatible if they belong to different classes, but bidirectionally incompatible if they are affiliated with the same class. Secondly, we assume that pollen is highly abundant and hence each diploid maternal plant has an unlimited number of fertilization attempts by incoming non-self pollen.

Similar to the full model, consider a population of $N$ diploid individuals. Each diploid is composed of two distinct haplotypes, where each haplotype belongs to one of $l$ classes. First set some notation. A haplotype class is represented by a number in $[\ell] := \{1, 2, \ldots, \ell\}$. A diploid individual is represented by a pair $(i, j)$ for some $i < j \in [\ell]$. We denote $X_t(i, j)$ for the number of diploids whose haplotypes belong to classes $i$ and $j$ (where $i < j$) at time $t$. For the sake of convenience, we write $X_t(i,j) = X_t(j,i)$ for $i > j$ in $[\ell]$, so that the order of indices does not matter. Write $X_t(i) := \sum_{k \neq i} X_t(i,k)$ for the number of individuals with one haplotype in class $i$. Lack of self-compatibility translates into the assumption that $X_t(i,i) = 0$ (no homozygotes) for all $i$ and $t$. The total fixed size of the population is given by $N = \sum_i \sum_{j>i} X_t(i,j)$, which is fixed at all times $t$. Accordingly, the total number of haplotypes is double, $\sum_i X_t(i) = 2N$.

At each time-step, we produce $N$ offspring by repeatedly selecting a diploid maternal plant uniformly from the previous generation's population and then selecting a compatible haplotype as sire, uniformly at random from all compatible ones. Thus, in the next generation a haplotype of type $i$ could be produced either as an offspring of a type $i$ ovule or type $i$ pollen, but not both. The probability of the former is simply the population proportion of type $i$ ovules $\frac{X_t(i)}{2N}$, because the ovules have an unlimited number of attempts to find partners and hence their fertilization is guaranteed. The probability of the latter is the frequency of diploid maternal plants compatible with type $i$ pollen

$$\sum_{\substack{k_1,k_2 \in [\ell] \setminus \{i\} \\ k_1 < k_2}} \frac{X_t(k_1,k_2)}{N},$$

multiplied by the proportion of type $i$ pollen out of all pollen compatible with that maternal plant

$$\frac{X_t(i)}{2N - X_t(k_1) - X_t(k_2)}.$$

Rearranging, this probability amounts to

$$\frac{X_t(i)}{N} \sum_{\substack{k_1,k_2 \in [\ell] \setminus \{i\} \\ k_1 < k_2}} \frac{X_t(k_1,k_2)}{2N - X_t(k_1) - X_t(k_2)}. \tag{1}$$

To simplify the analysis we concern ourselves only with the expected change in the ratio of the number of haplotypes of each class between generations. We therefore define $\sigma_t(i) := X_{t+1}(i)/X_t(i)$, the between-generations reproductive ratio of haplotype $i$ at time $t$. As $N$ offspring are produced every generation, we should add up both ovule and pollen (Eq. (1)) contributions to type $i$ offspring and multiply by $N$, so that

$$\begin{aligned} \mathbb{E}[\sigma_t(i) \mid X_t] &= \frac{1}{2} + \sum_{\substack{k_1,k_2 \in [\ell] \setminus \{i\} \\ k_1 < k_2}} \frac{X_t(k_1,k_2)}{2N - X_t(k_1) - X_t(k_2)} \\ &= \frac{1}{2} + \frac{1}{2} \sum_{\substack{k_1,k_2 \in [\ell] \setminus \{i\} \\ k_1 \neq k_2}} \frac{X_t(k_1,k_2)}{2N - X_t(k_1) - X_t(k_2)}. \end{aligned} \tag{2}$$

We turn to show that $X_t(i) \equiv \frac{2N}{\ell}$ is the only stable equilibrium of this equation. To see that it is indeed an equilibrium, we use Eq. (2) to compute:

$$\mathbb{E}\left[\sigma_t(i) \mid X_t(1) = \cdots = X_t(\ell) = \tfrac{2N}{\ell}\right] = \frac{1}{2} + \frac{N - \frac{2N}{\ell}}{2N - 2 \cdot \frac{2N}{\ell}} = 1. \tag{3}$$

As the expected reproductive rate of every haplotype type, without loss of generality, is 1, the state of equal proportions for all haplotypes is indeed an equilibrium point of the model. In order to establish uniqueness of this equilibrium, we study the extrema of $\mathbb{E}[\sigma_t(i) \mid X_t]$. Without loss of generality, we may omit the index $t$ and consider only the class $i = 1$. Here and in what follows we think only of $X = \{X(i,j)\}_{1<i<j\leqslant\ell}$ as a set of unknown variables, whereas $\{X(1,j)\}_{1<j\leqslant\ell}$ are fixed numbers. By Eq. (2), we are looking for the extrema of the function

$$f(X) = \sum_{\substack{k_1,k_2\in[\ell]\setminus\{1\} \\ k_1\neq k_2}} \frac{X(k_1,k_2)}{2\sum\limits_{j,m\notin\{k_1,k_2\}} X(j,m) + \sum\limits_{m\neq k_2} X(k_1,m) + \sum\limits_{j\neq k_1} X(j,k_2)}. \tag{4}$$

under the constraints $\sum_{i,j} X(i,j) = N$ and $X(i,j) \geqslant 0$ for all $i,j \in [\ell]$. By Lagrange's multipliers method, if $X^*$ is an extremal point under these constraints, then there is some $\lambda \in \mathbb{R}$ such that

$$2 \leqslant k_1 < k_2 \leqslant \ell \Rightarrow \frac{\partial f}{\partial X(k_1,k_2)}(X^*) = \lambda. \tag{5}$$

A direct computation yields

$$\begin{aligned} \frac{\partial f}{\partial\{X(k_1,k_2)\}} &= \frac{1}{2N - X(k_1) - X(k_2)} - \sum_{j,m\in[\ell]\setminus\{k_1,k_2,1\}} \frac{2X(j,m)}{(2N - X(j) - X(m))^2} \\ &- \sum_{m\in[\ell]\setminus\{k_1,k_2,1\}} \frac{X(k_1,m)}{(2N - X(k_1) - X(m))^2} - \sum_{j\in[\ell]\setminus\{k_1,k_2,1\}} \frac{X(j,k_2)}{(2N - X(k_2) - X(j))^2} \end{aligned} \tag{6}$$

To ease the notation, denote

$$A(j,m) = \frac{X(j,m)}{(2N - X(j) - X(m))^2} \tag{7}$$

and

$$A(j) = \sum_{m\neq 1} A(j,m), \quad S = \sum_{j\neq 1}\sum_{m\neq 1} A(j,m).$$

Then

$$\begin{aligned} \frac{\partial f}{\partial\{X(k_1,k_2)\}} =& \frac{1}{2N - X(k_1) - X(k_2)} \\ & - \left( S - A(k_1,k_2) + \sum_{j,m\in[\ell]\setminus\{k_1,k_2,1\}} A(j,m) \right). \end{aligned}$$

By Eq. (5), we deduce that $\frac{\partial f}{\partial\{X(k_1,k_2)\}} = \frac{\partial f}{\partial\{X(k_1,k_3)\}}$ for all different $k_1$, $k_2$ and $k_3$ in $\{2,\ldots,\ell\}$, so that, by the last equation we have:

$$\begin{aligned} &\frac{1}{2N - X(k_1) - X(k_2)} - \frac{1}{2N - X(k_1) - X(k_3)} + A(k_1,k_2) - A(k_1,k_3) \\ &= \sum_{j,m\in[\ell]\setminus\{1,k_1,k_2\}} A(j,m) - \sum_{j,m\in[\ell]\setminus\{1,k_1,k_3\}} A(j,m) \\ &= A(k_3) - A(k_2) - A(k_3,k_1) + A(k_2,k_1). \end{aligned}$$

All in all, we arrive at

$$\frac{1}{2N - X(k_1) - X(k_2)} - \frac{1}{2N - X(k_1) - X(k_3)} = A(k_3) - A(k_2). \tag{8}$$

Since the right-hand-side is independent of $k_1$, we deduce that also the left-hand-side must be so. For $\ell \geqslant 5$, this is possible only if $\{X(k)\}_{k \neq 1}$ are all equal.[1] Indeed, assume for the sake of contradiction that $X(k) \neq X(k')$ for some $k, k' \in [\ell] \backslash \{1\}$, then by choosing $k_2, k_3 \in [\ell] \backslash \{1, k, k'\}$ and writing Eq. (8) once with $k_1 = k$ and once with $k_1 = k'$, we shall get a contradiction. We deduce that, given $X(1)$, the minimum of the expectation of $\sigma(1)$ is attained when

$$X(k) = \frac{2N - X(1)}{\ell - 1}, \quad \forall k \in [\ell] \backslash \{1\}. \tag{9}$$

Observe that under the conditions of Eq. (9), the value of the function $f$ (defined in Eq. (4)) depends only on $X_t(1), \ldots, X_t(\ell)$:

$$f(X_{\text{eq}}) = \frac{N - X(1)}{2N - 2 \cdot \frac{2N - X(1)}{\ell - 1}}.$$

This is clearly strictly monotone decreasing in the value of $X(1)$. By Eq. (3), we deduce that

$$\mathbb{E}\left[\sigma_t(1) \mid X_t(1) < \tfrac{2N}{\ell}\right] > 1. \tag{10}$$

A similar inequality holds for any $i \in [\ell]$. We deduce that $X_t(i) \equiv 2N/\ell$ is a unique attractive equilibrium. Of course, if any of the classes is extinct the system will drift towards a new stable equilibrium of perfect balance between the remaining classes, but this is not an attractor equilibrium, as in its vicinity there are points where the system is repelled from it.

To obtain the approximated behaviour of the process for large populations, we can consider the more streamlined Moran model, in which at every time step, a single diploid is produced, replacing a random diploid of the previous population. We denote the number of copies of a haplotype of class $i$ in this process by $X'_t(i)$. Observe that time here is scaled down by a factor of $N$. Denoting $\sigma'_t(i) = X'_{t+1}/X'_t$, we observe that $X'_t(i)$ is modified at each time step by $B_i(t) - D_i(t)$ where $B_i \sim \text{Ber}\left(\frac{X'_t(i)\sigma'_t(i)}{N}\right)$ and $D_i \sim \text{Ber}\left(\frac{X'_t(i)}{N}\right)$. Hence, by the Azuma–Hoeffding inequality, which states that the sum of independent Bernoulli random variables is exponentially concentrated around its mean, we deduce that $X'_t(i)$ is exponentially concentrated about $\sum_{s=0}^{t} \frac{X'_s(i)(\sigma'_s(i)-1)}{N}$. In particular, we deduce that the probability of the process $X'_t(i)$ hitting 0, starting an excursion from $X'_0(i) \geqslant \frac{cN}{\ell}$ for a given $c > 0$, is exponentially unlikely in $N$. A similar argument implies the same conclusion for $X_t(i)$. We conclude that for large populations, extinctions of sizable classes due to genetic drift should be exceedingly rare.

---

[1] The cases $\ell = 3$ and $\ell = 4$ are analyzed directly. For $\ell = 4$ the unique equilibrium point is found by solving the linear system of equations in three variables $X(2,3)$, $X(2,4)$, $X(3,4)$ given by the constraint $\sum_{i<j} X(i,j) = N$, Eq. (8) taken with $(k_1, k_2, k_3) = (2, 3, 4)$ and later with $(k_1, k_2, k_3) = (3, 2, 4)$; by symmetry, the solution obeying $X(2) = X(3) = X(4)$ is the unique solution and we proceed as with $\ell \geqslant 5$. For $\ell = 3$ Eq. (2) yields $\mathbb{E}[\sigma_t(1)|X_t] = \frac{N}{X(1)} - \frac{1}{2}$, which is monotone decreasing in $X(1)$, arriving immediately at Eq. (10).

#### The survival time of the endangered sub-class awaiting a rescue mutation during the class split process

In all split trajectories of a given class (hereby class 1), the second mutation grants a certain sub-class ('driver', denoted by $1_+$) compatibility as sire with another sub-class ('endangered', denoted by $1_-$). This reproductive benefit triggers the expansion of the driver sub-class over both the endangered subclass and the remainder of the class ('neutral', denoted by $1_*$). To complete the split, a third mutation should occur in the endangered sub-class, whose numbers are in decline. Hence, this mutation must occur within a limited time window before that haplotype is lost. To analyze the time window affording this mutation, and, more importantly, the total number of individuals over all generations in this time window (measuring the mutation opportunities for the final mutation in the splitting process), we study first the growth of the driver class $1_+$ and then the decline of the endangered class $1_-$. To study this, we update the set of haplotype classes to be $L = \{1_-, 1_*, 1_+, 2, \ldots, \ell\}$. Class $1_-$ is incompatible as sire with dams of types $1_*$ and $1_+$; class $1_*$ is incompatible as sire with dams of types $1_-$ and $1_+$; while class $1_+$ is incompatible as sire only with dams of type $1_*$. All other between-class mating is possible, and the evolution of the model is just as before under these constraints. Thus, denoting $L_1 = \{1_-, 1_*, 1_+\}$, the expected reproduction ratio $\sigma_t(i) = X_{t+1}(i)/X_t(i)$ is still given by Eq. (2) for $i \notin L_1$ (where $[\ell]$ is replaced by $L$), while for $i \in L_1$ it is replaced by

$$\mathbb{E}[\sigma_t(i) \mid X_t] = \frac{1}{2} + \frac{1}{2} \sum_{\substack{k_1,k_2 \in L\setminus J_i \\ k_1 \neq k_2}} \frac{X_t(k_1,k_2)}{2N - X_t^*(k_1) - X_t^*(k_2)}, \tag{11}$$

where $J_{1_-} = J_{1_*} = L_1$ and $J_{1_+} = \{1_+, 1_*\}$ and $X_t^*(k) = \sum_{j \in J_k} X_t(j)$ for all $k \in L_1$ while $X_t^*(k) = X_t(k)$ for all $k \notin L_1$. In particular, we observe that $\mathbb{E}[\sigma_t(0) \mid X_t] = \mathbb{E}[\sigma_t(-1) \mid X_t]$, which leads to

$$\frac{\mathbb{E}\left[X_{t+1}(1_*) \,\middle|\, X_t\right]}{\mathbb{E}\left[X_{t+1}(1_-) \,\middle|\, X_t\right]} = \frac{X_t(1_*)}{X_t(1_-)}.$$

We therefore conclude that, on average, the proportion

$$\theta_t = \frac{X_t(1_-)}{X_t(1_*) + X_t(1_-)} \tag{12}$$

remains constant.

#### Mutation trajectories starting with an RNase mutation

We begin by analysis of the split trajectories initiated by an RNase mutation. As discussed above, In these trajectories, for the parameters of our simulation, the occurrence of a third mutation is rather commonplace. This merits analysis of the mean evolution of each class for the purpose of estimating the typical number of mutation opportunities.

With Eq. (12) in mind, we firstly assume, as a simplification, that $\theta_t \equiv \theta \in [0, 1]$ is fixed for all $t$ in our evolutionary trajectory. We start with the analysis of the class driving the endangered sub-class to extinction (in our notation, class $1_+$). Assuming the initial counts of individuals in the special classes of $L_1$ are given, we can conclude, by similar arguments to those leading to Eq. (9), that all other classes are balanced among themselves, on average. The total population of the classes of $L_1$ are not expected to be equal to the remaining classes, but rather somewhat larger, as class $1_+$ has more mating partners than elements of other classes. However, this is not a large increase (see Fig. S2), bounded from above by the size of $X_{1_-}$, and as it affects the size of the total class 1 much more than it affects the relative proportion of its different components, as long as $X_{1_-}$ is significant. We thus simplify the analysis by forcing $X_t(i) = \frac{2N}{\ell}$ for all $i \in L \backslash L_1$. We expect this analysis to be reliable for all sub-classes at least until the variant $X_{1_+}$ becomes prevalent, namely until the first time $t_{\text{prev}}$ for which $X_{t_{\text{prev}}}(1_+) = \frac{N}{\ell}$. This approximation, however, is invalid for the study of the final decline of $X_{1_-}$ after $t_{\text{prev}}$, as the pressure that it puts on $X_{1_-}$ to vanish is too strong. Hence, in order to analyze the evolution of $X_{1_-}$ from $t_{\text{prev}}$ and on, we shall later take the recovered solution for the evolution of $X_{1_+}$ as the assumption, and allow the size of the entire $X_1$ class to vary.

Under $X_t(i) = \frac{2N}{\ell}$ for all $i \geqslant 2$, we have, by Eq. (11),

$$\begin{aligned}
&\mathbb{E}\left[\frac{X_{t+1}(1_+)}{X_t(1_+)} \,\middle|\, X_t(2) = \cdots = X_t(\ell) = \frac{2N}{\ell}\right] \\
&= \frac{1}{2} + \frac{\frac{1}{2}\sum_{k=2}^{\ell} X_t(k)}{2N - \frac{4N}{\ell}} + \frac{\frac{1}{2}X_t(1_-)}{2N - \frac{2N}{\ell} - (\frac{2N}{\ell} - X_t(1_-))} \\
&= \frac{1}{2} + \frac{N - \frac{2N}{\ell}}{2N(1-\frac{2}{\ell})} + \frac{\theta\left(\frac{N}{\ell} - \frac{1}{2}X_t(1_+)\right)}{2N(1-\frac{2}{\ell}) + \theta\left(\frac{2N}{\ell} - X_t(1_+)\right)} \\
&\approx \frac{1}{2} + \frac{(1-\frac{2}{\ell})N + \theta\left(\frac{N}{\ell} - \frac{1}{2}X_t(1_+)\right)}{2N(1-\frac{2}{\ell})}
\end{aligned}$$

Writing $X_t^+ = X_t(1_+)$ for short, and using the identity $\frac{X_{t+1}}{X_t} = \frac{X_{t+1}-X_t}{X_t} + 1$, we approximate the dynamics by the mean continuous time equation:

$$\frac{dX_t^+/dt}{X_t^+} = \frac{(1-\frac{2}{\ell})N + \theta\left(\frac{1}{\ell}N - \frac{1}{2}X_t^+\right)}{2N(1-\frac{2}{\ell})} - \frac{1}{2} = \frac{\frac{\theta}{\ell}N - \frac{\theta}{2}X_t^+}{N(1-\frac{2}{\ell})}. \tag{13}$$

Following these approximations, the solution is readily found by separation of variables to be

$$X_t(1_+) = \frac{2N}{\ell}\left(1 - \frac{1}{c\exp\left(\frac{\theta}{\ell-2}t\right) + 1}\right), \tag{14}$$

where

$$c = \frac{\ell X_0(1_+)}{2N - \ell X_0(1_+)} \tag{15}$$

is a constant determined by the initial conditions. In particular we observe that the size of class $1_+$ increases asymptotically to $2N/\ell$, which is the size of all other classes in $\{2, \ldots, \ell\}$.

We turn to estimate the evolution of the endangered subclass $1_-$ in this time window. The solution in Eq. (14), together with Eq. (12) and the balanced assumption $X(1_+) + X(1_*) + X(1_-) = \frac{2N}{\ell}$, yield the following solution for $X(1_-)$:

$$X_t(1_-) = \theta\left(\frac{2N}{\ell} - X_t(1_+)\right) = \frac{\theta\frac{2N}{\ell}}{c\exp\left(\frac{\theta}{\ell-2}t\right) + 1}. \tag{16}$$

This solution is valid until $t_{\text{prev}}$, after which which class $1_+$ becomes prevalent, that is, $X_{t_{\text{prev}}}(1_+) =$

$\frac{N}{\ell}$. By Eq. (14) it obeys $\exp\left(\frac{\theta}{\ell-2}t_{\text{prev}}\right) = \frac{1}{c}$, or equivalently

$$t_{\text{prev}} = \frac{\ell-1}{\theta}\ln\left(\frac{N-\ell X_0^+}{\ell X_0^+}\right). \tag{17}$$

As $X_t(1_+)$ tends to $\frac{2N}{\ell}$, though, it becomes less and less valid. This is due to the fact that the pressure on $X_{1_+}$ to grow to size $\frac{2N}{\ell}$ is much stronger than the pressure on $X_{1_-}$ to die out. From this point and on, we thus abandon our assumption that $X_1 = \frac{2N}{\ell}$ and instead assume the validity of the evolution of $X_{1_+}$ which we have recovered earlier, and use it to recover the evolution of $X_{1_-}$ separately. To do so we write the evolution of $X_{1_-}$ according to Eq. (11):

$$\begin{aligned}\mathbb{E}\left[\frac{X_{t+1}(1_-)}{X_t(1_-)}\,\middle|\, X_t(2)=\cdots=X_t(\ell)\right] &= \frac{1}{2}+\frac{1}{2}\cdot\frac{\sum_{k=2}^{\ell}X_t(k)}{2N-2\frac{2N}{\ell}}\\ &= \frac{1}{2}+\frac{N-X_t(1_+)-(X_t(1_-)+X_t(1_*))}{2N(1-\frac{2}{\ell})}\\ &= \frac{1}{2}+\frac{N-\frac{N}{\ell}\left(1-\frac{1}{c\exp(\frac{\theta}{\ell-2}t)+1}\right)-\frac{1}{\theta}X_t(1_-)}{2N(1-\frac{2}{\ell})}\end{aligned}$$

Similarly to Eq. (13), we move to the continuous time differential equation

$$\frac{X_t'(1_-)}{X_t(1_-)} = \frac{N-\frac{N}{\ell}\left(1-\frac{1}{c\exp(\frac{\theta}{\ell-2}t)+1}\right)-\frac{1}{\theta}X_t(1_-)}{2N(1-\frac{2}{\ell})}-\frac{1}{2} = \frac{\frac{1}{c\exp(\frac{\theta}{\ell-2}t)+1}-\frac{1}{\theta}X_t(1_-)}{2N(1-\frac{2}{\ell})}.$$

After a change of variables this equation can be solved by separation of variables, which yields

$$X_t(1_-) = 2\theta N(1-\tfrac{2}{\ell})\exp\left(\frac{f(t)}{2N(1-\frac{2}{\ell})}\right)\frac{1}{\int\exp\left(\frac{f(t)}{2N(1-\frac{2}{\ell})}\right)dt}, \tag{18}$$

where

$$f(t) = \int\frac{dt}{c\exp(\frac{\theta}{\ell-2}t)+1} = \ln\left(1-\frac{1}{1+c\exp\left(\frac{\theta}{\ell-2}t\right)}\right) \approx \frac{1}{1+c\exp\left(\frac{\theta}{\ell-2}t\right)}.$$

The indefinite integral in Eq. (18) is

$$\begin{aligned}\int\exp\left(\frac{f(t)}{2N(1-\frac{2}{\ell})}\right)dt &= \int\left(1-\frac{1}{1+c\exp(\frac{\theta}{\ell-2}t)}\right)^{\frac{1}{2N(1-\frac{2}{\ell})}}dt\\ &\approx \int\left(1-\frac{1}{\left(1+c\exp(\frac{\theta}{\ell-2}t)\right)\cdot 2N\left(1-\frac{2}{\ell}\right)}\right)dt\\ &\approx (t-t_{\text{prev}})+\frac{1-\exp(-\frac{\theta}{\ell-2}t)}{\frac{c\theta}{\ell-2}\cdot 2N(1-\frac{2}{\ell})}+C\\ &\approx (t-t_{\text{prev}})+\frac{\ell}{2N\cdot c\theta}\left(1-\exp\left(-\frac{\theta}{\ell-2}t\right)\right)+C,\end{aligned}$$

where $C$ is a constant that should be chosen according to the initial conditions. Plugging this back into Eq. (18), and taking into account the initial conditions $X_{t_{\text{prev}}}(1_-) = \frac{\theta N}{\ell}$ (clf. Eq. (16) and Eq. (17)), we see that, for $t > t_{\text{prev}}$,

$$X_t(1_-) \approx \frac{2\theta N(1-\frac{2}{\ell})}{(t-t_{\text{prev}})+2\ell-4}, \quad t > t_{\text{prev}}. \tag{19}$$

The time $t_{\text{ext}}$ at which class $1_-$ is nearly extinct, that is, $X_{t_{\text{ext}}}(1_-) = X^-_{\text{ext}}$, could now be approximated by

$$t_{\text{ext}} = t_{\text{prev}} + \frac{2\theta(1-\frac{2}{\ell})}{X^-_{\text{ext}}}N - 2\ell + 4 \tag{20}$$

At last, we can calculate the cumulative number of individuals in the endangered class in all generations, from the time of the second mutation ($t = 0$) and until it is nearly extinct ($t = t_{\text{ext}}$), which together are proportional to the number of opportunities to gain the rescue mutation and evade extinction. Up to time $t_{\text{prev}}$ we use Eq. (16) and get

$$\begin{aligned}
\int_0^{t_{\text{prev}}} X_t(1_-)dt &= \int_0^{t_{\text{prev}}} \frac{\theta\frac{2N}{\ell}}{ce^{\frac{\theta}{\ell-2}t}+1}dt \\
&= 2\frac{\theta N}{\ell}\cdot\frac{\ell-2}{\theta}\ln\left(\frac{ce^{\frac{\theta}{\ell-2}t}}{ce^{\frac{\theta}{\ell-2}t}+1}\right)\Bigg|_0^{t_{\text{prev}}} \\
&= 2\frac{\ell-2}{\ell}N\left\{\ln\left(\frac{1}{2}\right)-\ln\left(\frac{c}{c+1}\right)\right\} \\
&= 2\frac{\ell-2}{\ell}N\ln\left(\frac{N}{\ell X_0^+}\right)
\end{aligned}$$

After time $t_{\text{prev}}$ we use Eq. (19) and Eq. (20):

$$\begin{aligned}
\int_{t_{\text{prev}}}^{t_{\text{ext}}} X_t(1_-)dt &\approx 2\theta(1-\tfrac{2}{\ell})N\left(\ln(t_{\text{ext}}-t_{\text{prev}}+2\ell-4)-\ln(2\ell-4)\right) \\
&= 2\theta(1-\tfrac{2}{\ell})N\ \ln\left(\frac{\theta}{\ell X^-_{\text{ext}}}N\right)
\end{aligned}$$

Thus the total number of individuals in the endangered class is

$$2(1-\tfrac{2}{\ell})N\ln\left(\frac{N}{\ell X_0^+}\right) + 2\theta(1-\tfrac{2}{\ell})N\ \ln\left(\frac{\theta N}{\ell X^-_{\text{ext}}}\right).$$

Observe that the probability that the third ('rescue') mutation occurs before the $1_+$ sub-class becomes prevalent is comparable to the probability that it occurs after this time, for low mutation rates.

Although this analysis is approximate, its validity should increase as the number of individuals in each class and in the endangered sub-class increases. In particular we draw from this the conclusion that the asymptotics of typical time to extinction when a mutation does not occur is $\Theta(N)$, while the time until the driver subclass $1_+$ become prevalent is only of order $\Theta(\ln N)$. The number of mutation opportunities offered for the third mutation by this process is $\Theta(N\ln(N/\ell))$.

Plugging in the numerical values of the simulation of figure S1**(a)**, namely $\ell = 8$ classes for

$N = 500$ individuals with $\theta = 0.44$, and starting from $X_+(0) = 11$, we obtain about 2468 mutation opportunities (cp. $n_\mu = 3768$ in simulation). For figure S1(**b**), with $N = 500$, $\ell = 9$ and $\theta = 0.17$, and starting from $X_+(0) = 13$ we obtain 1479 mutation opportunities (cp. $n_\mu = 1643$ in simulation).

#### Remarks concerning the split trajectory starting with an SLF mutation

Snce in the third split trajectory the SLF mutation occurs first and the RNase mutation compatible with that SLF is second, an analogous analysis should begin with only a single individual in the endangered sub-class 1-. It would therefore be meaningless to assume that $X_t(1_-)/X_t(1_*)$ is constant throughout the simulation, as it would make no sense to look at a typical evolution. Even in a setting of genetic drift, martingale arguments indicate that the probability of class $1_-$ becoming prevalent is of order $\Theta(\frac{\ell}{2N})$. Hence it is only the promiscuity of compatibility of RNase mutations to existing SLFs that allows this trajectory to occur in significant numbers. As $\frac{N}{L}$ grows we expect this trajectory to become rarer and rarer.

Table S1: AA frequencies and biochemical category by UniProt database

| AAs | percentage | bio-chemical classification |
|---|---|---|
| F(Phe) | 3.86 | hydrophobic |
| W(Trp) | 1.09 | hydrophobic |
| M(Met) | 2.41 | hydrophobic |
| L(Leu) | 9.65 | hydrophobic |
| I(Ile) | 5.93 | hydrophobic |
| V(Val) | 6.86 | hydrophobic |
| G(Gly) | 7.08 | hydrophobic |
| P(Pro) | 4.72 | hydrophobic |
| A(Ala) | 8.26 | hydrophobic |
| C(Cys) | 1.37 | neutral polarized |
| Y(Tyr) | 2.92 | neutral polarized |
| T(Thr) | 5.35 | neutral polarized |
| S(Ser) | 6.60 | neutral polarized |
| N(Asn) | 4.06 | neutral polarized |
| H(His) | 2.27 | neutral polarized |
| Q(Gln) | 3.93 | neutral polarized |
| E(Glu) | 6.74 | Negative charged |
| D(Asp) | 5.46 | Negative charged |
| R(Arg) | 5.53 | Positive charged |
| K(Lys) | 5.82 | Positive charged |

### Supplementary figures

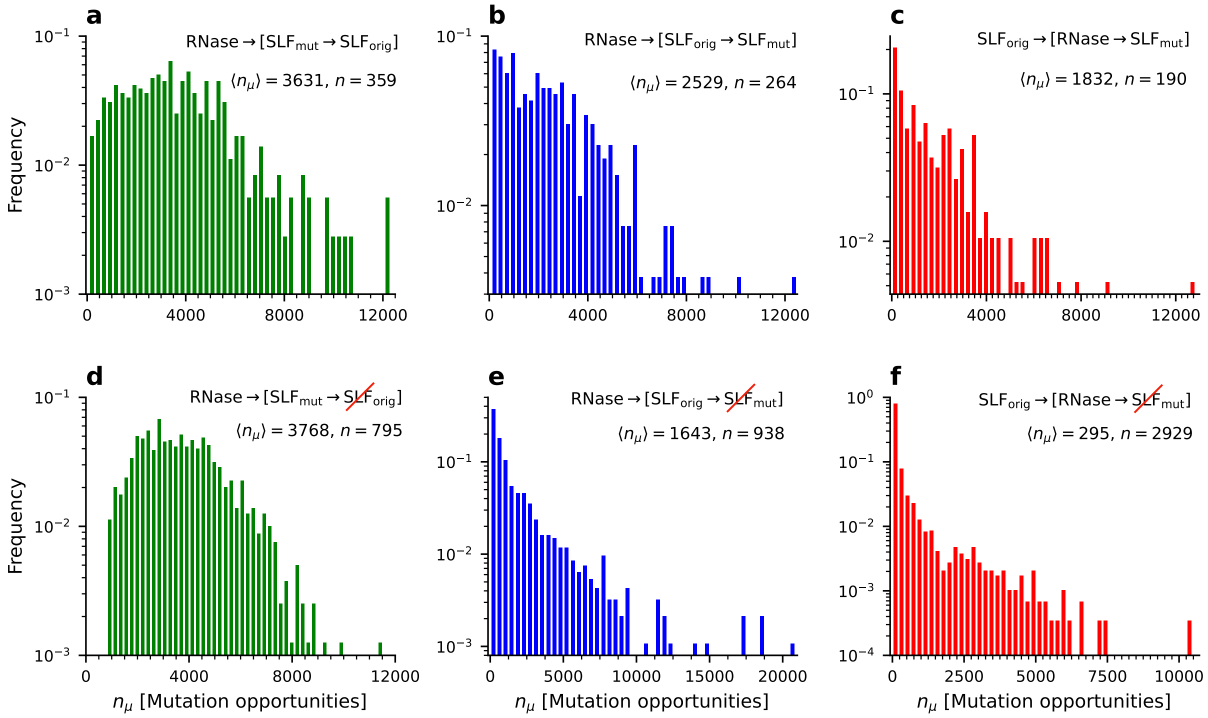


Figure S1: **Distribution of waiting times for split completion**. In all three split trajectories, the third mutation should occur on a selectively disadvantageous sub-class ('endangered'), whose population is under decline. Hence, to complete the split, there is a limited time in which the third ('rescue') mutation could occur before that sub-class vanishes. In **(a-c)** we show the distributions of 'mutation opportunities' $n_\mu$, in all completed splits as observed in our simulations, dissected by their trajectory type. We define mutation opportunities as the cumulative copy number $X(t)$ of the sub-class that should receive the third mutation, between the occurrences of the second mutation $T_2$ and third mutation $T_3$, $n_\mu = \sum_{t=T_2}^{T_3} X(t)$. **(d-f)** To quantitate the feasible time in which the third mutation could occur before the extinction of the endangered sub-class, we chose the three split examples illustrated in Fig. 7 as test cases. For each of them, we took the population state a few generations after the second mutation as the starting point. From that point, we ran multiple forward simulations of selection and drift only, where mutations were disabled to avoid further changes. In each run, we assessed the time to extinction of the endangered, conditioned on the survival of the haplotype carrying the second mutation, which drives its extinction ('driver'). Similar to (a-c) we show here the distributions of mutation opportunities, over these multiple stochastic repeats, with the only difference that now they are calculated between the second mutation and the extinction time $T_\infty$ of the endangered sub-class, $n_\mu = \sum_{t=T_2}^{T_\infty} X(t)$. Note that while (a-c) show distributions over the many different split events observed in simulation, (d-f) focus on three particular cases. $\langle \Delta n_\mu \rangle$ is the distribution mean in each case. $n$ is the number of realizations used to form the distribution.

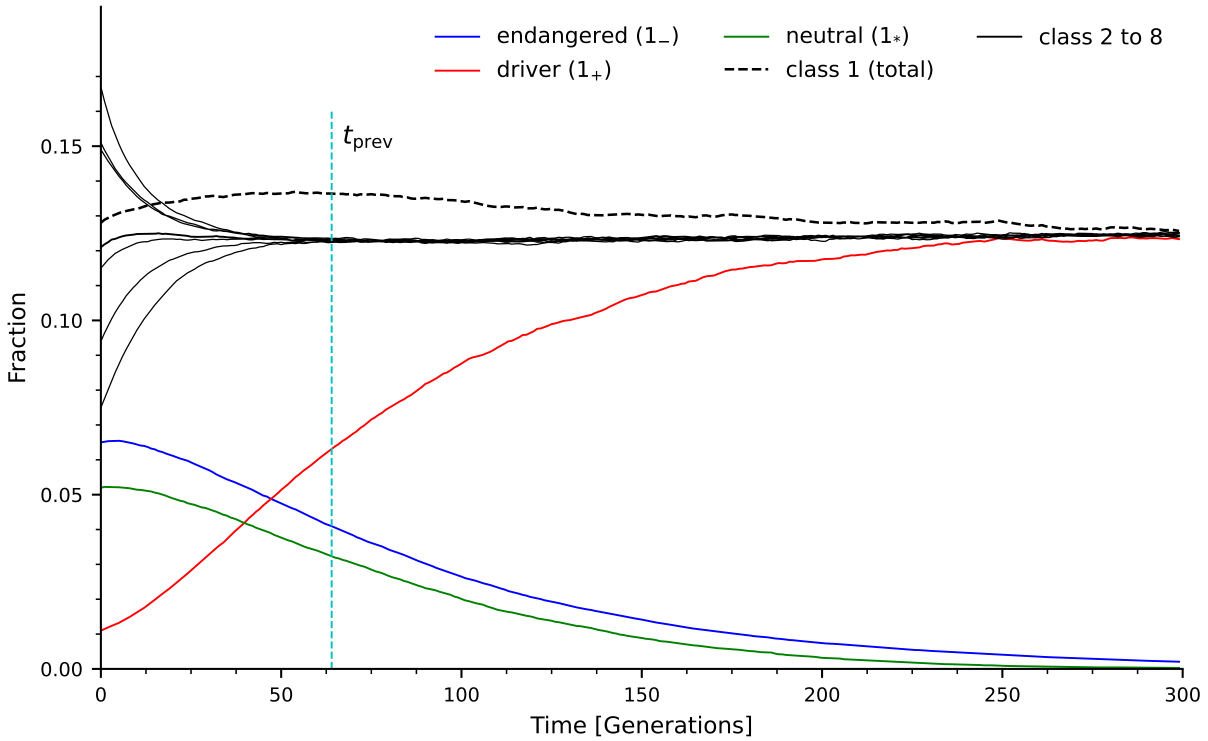


Figure S2: **Dynamics of the endangered, neutral, and driver sub-classes in the absence of mutations**. Using the initial conditions of the first split trajectory shown in Fig. 7 we ran stochastic simulations of the selection-drift dynamics of all existing population classes when no additional mutations are enabled. The driver sub-class ($1_+$, red) is advantageous as it is compatible as a sire with the endangered sub-class ($1_-$, blue) besides all classes the endangered and neutral ($1_*$, green) are compatible with. Thus, the driver class causes the extinction of both the endangered and neutral sub-classes and takes over the entirety of class 1. All other classes reach the equilibrium point of equal proportions. Due to the absence of mutations, no additional classes or sub-classes can emerge. The results are averages over 1000 repeats of the stochastic simulation starting from the same initial conditions. In each simulation we condition on the survival of the driver and the endangered classes, otherwise the simulation is stopped. This leads to under-representation of the neutral (which could also vanish) in late times.

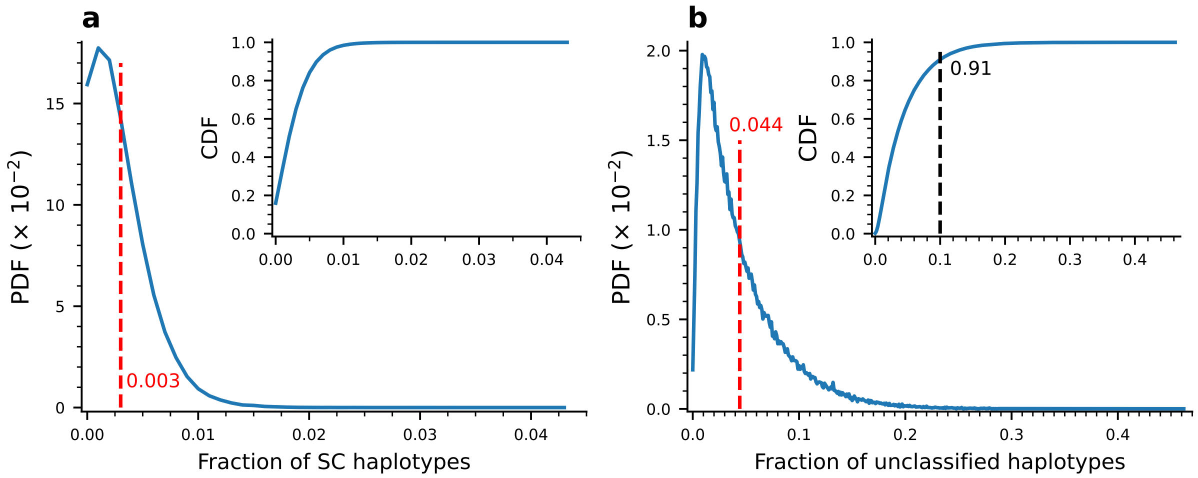


Figure S3: **The vast majority of the population is associated with any of the compatibility classes, $E_{\mathrm{th}} = -6$, $N = 500$.** This figure complements Fig. 5. **(a)** Distribution of the proportion of self-compatible haplotypes, and **(b)** Distribution of the proportion of unclassified haplotypes across multiple simulation instances. The red dashed vertical lines in both panels show the distribution mean. The insets in both panels show the cumulative distributions. In the inset, the black vertical dashed line shows that in 91% of the instances, at least 90% of the haplotypes are classified.

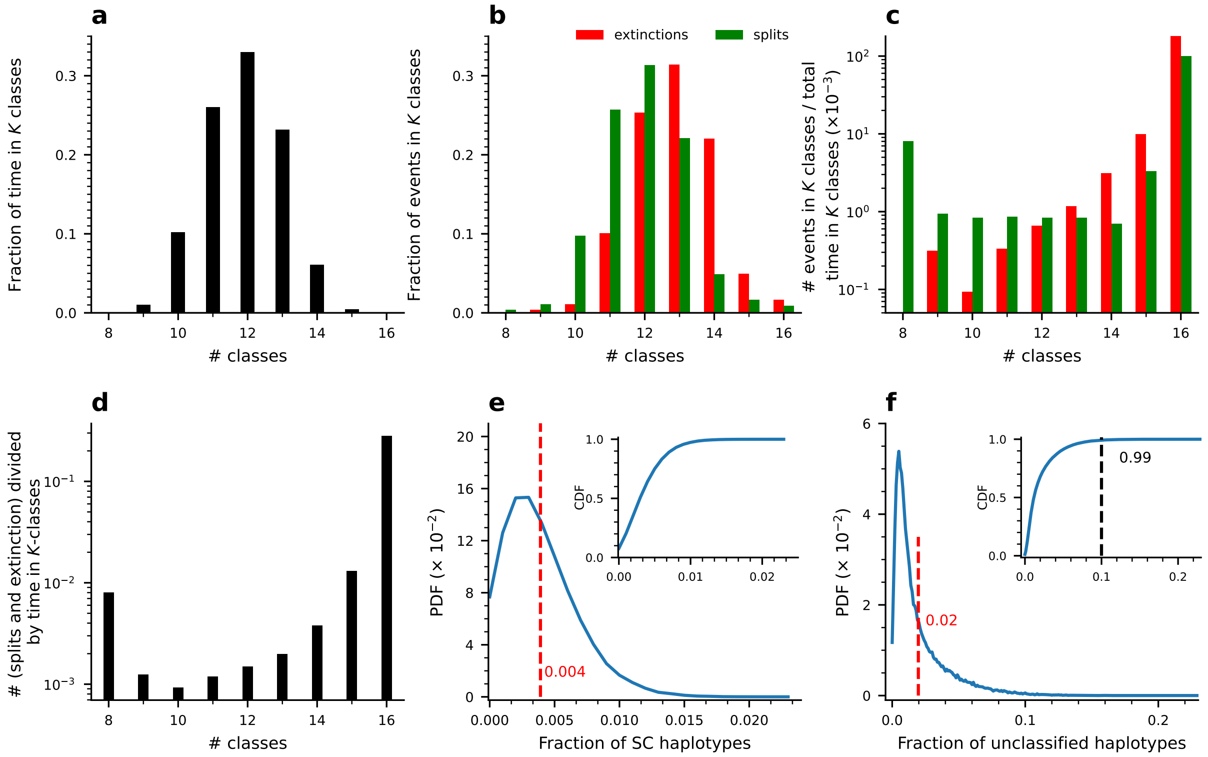


Figure S4: **The equilibrium number of classes increases with $E_{\mathbf{th}}$, but the model retains its qualitative behavior, simulation results with $E_{\mathbf{th}} = -4$, $N = 500$.** **(a)** The fraction of time spent under $K$-classes state. **(b)** The fraction of split (green) and extinction (red) events that occurred under each $K$-class state. **(c)** The extinction (split) rate – the number of extinction (red) and split (green) events under each $K$-classes state divided by the time spent at this state shown against $K$. (a-c) demonstrate a stable equilibrium, similar to Fig. 5, but with equilibrium value of $K = 12$. Comparison to Fig. 5 where $E_{\text{th}} = -6$ was used, shows that a higher $E_{\text{th}}$ value, which amounts to higher probability to interact at random, results in a higher number of classes, for the same population size. **(d)** Total event rate: sum of both split and extinction events under $K$ classes, divided by the time spent under $K$ classes (sum of the red and green bars as they appear in (c)). **(e)** Distribution of the proportion of self-compatible haplotypes, and **(f)** Distribution of the proportion of unclassified haplotypes across multiple simulation instances. The red dashed vertical lines in both panels show the distribution mean. The insets in both panels show the cumulative distributions. Results based on 19 independent runs, with a total of 871620 generations.

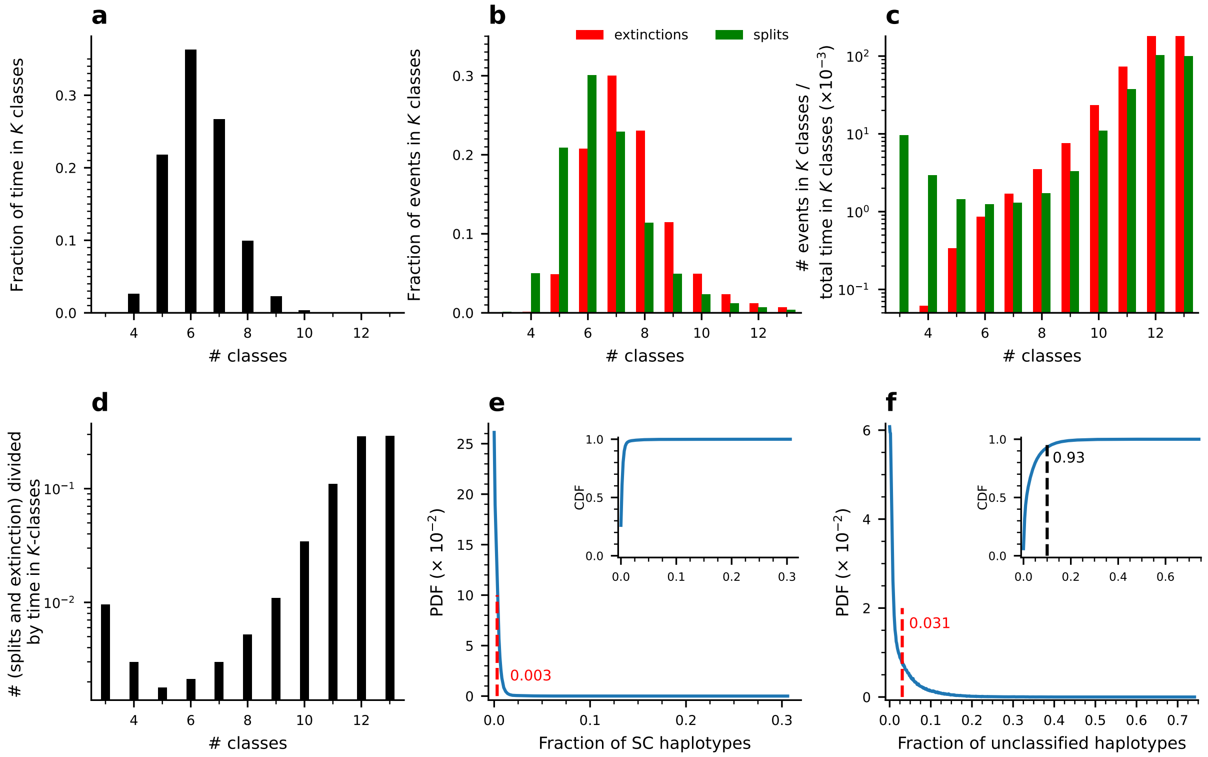


Figure S5: **The equilibrium number of classes decreases when $E_{th}$ decreases, but the model retains its qualitative behavior, simulation results for $E_{th} = -8$, $N = 500$.** **(a)** The fraction of time spent under $K$-classes state. **(b)** The fraction of split (green) and extinction (red) events that occurred under $K$-class state. **(c)** The extinction (split) rate – the number of extinction (red) and split (green) events in $K$-classes state divided by the time spent at this state – against $K$. (a-c) demonstrate a stable equilibrium, similar to Fig. 5, but a lower equilibrium value of $K = 6$. Comparison to Fig. 5 where $E_{th} = -6$ was used, shows that a lower $E_{th}$ value, which amounts to lower probability to interact at random, results in a lower number of classes, for the same population size. **(d)** Total event rate: sum of both split and extinction events under $K$ classes, divided by the time spent under $K$ classes (sum of the red and green bars as they appear in (c)). **(e)** Distribution of the proportion of self-compatible haplotypes, and **(f)** Distribution of the proportion of unclassified haplotypes across multiple simulation instances. The red dashed vertical lines in both panels show the distribution mean. The insets in both panels show the cumulative distributions. Results based on 59 independent runs, with a total of 3890920 generations.

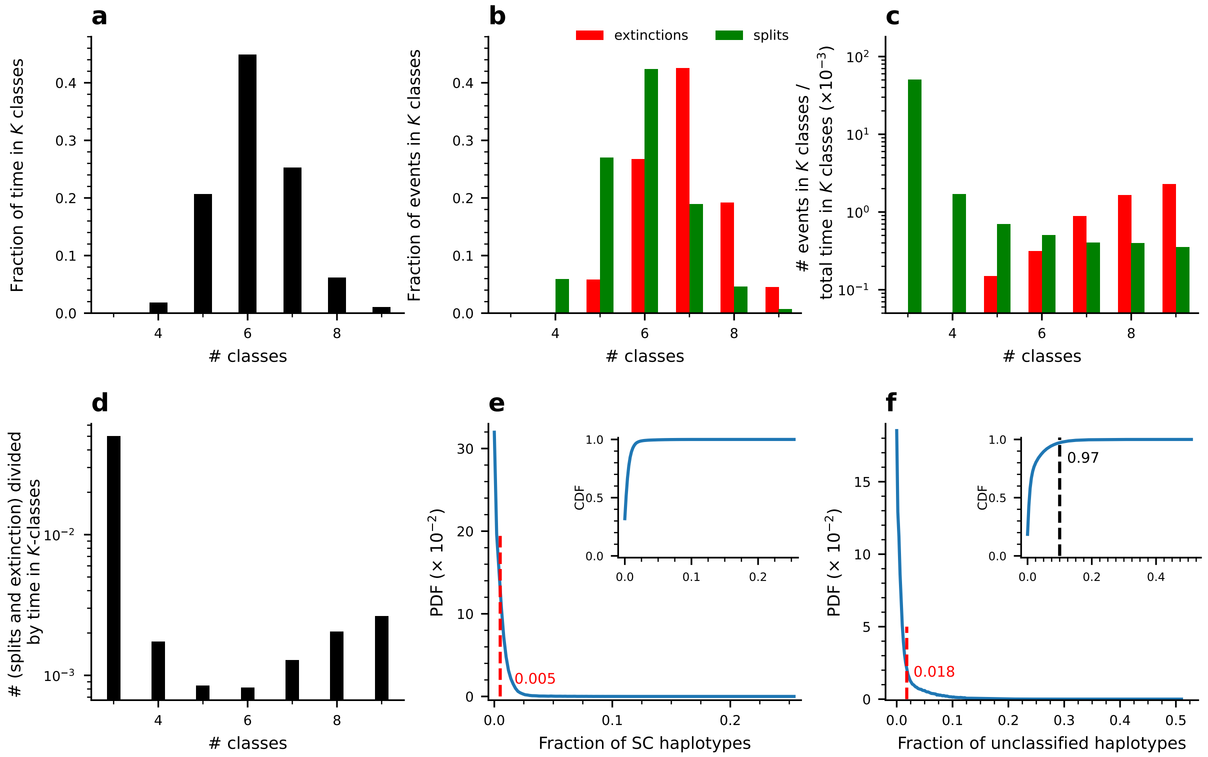


Figure S6: **The equilibrium number of classes decreases when the population size decreases, but the model retains its qualitative behavior, simulation results for $E_{\mathbf{th}} = -6$, and $N = 250$.** **(a)** The fraction of time spent under $K$-classes state. **(b)** The fraction of split (green) and extinction (red) events that occurred under $K$-class state. **(c)** The extinction (split) rate – the number of extinction (red) and split (green) events in $K$-classes state divided by the time spent at this state – against $K$. (a-c) demonstrate a stable equilibrium, similar to Fig. 5, but with a lower equilibrium value of $K = 6$. Comparison to Fig. 5 where $N = 500$ was used, shows that lower $N$ results in a lower number of classes, for the same energy threshold value, $E_{\mathrm{th}} = -6$. **(d)** Total event rate: sum of both split and extinction events under $K$ classes, divided by the time spent under $K$ classes (sum of the red and green bars as they appear in (c)). **(e)** Distribution of the proportion of self-compatible haplotypes, and **(f)** Distribution of the proportion of unclassified haplotypes across multiple simulation instances. The red dashed vertical lines in both panels show the distribution mean. The insets in both panels show the cumulative distributions. Results based on 25 independent runs, with a total of 1714440 generations.

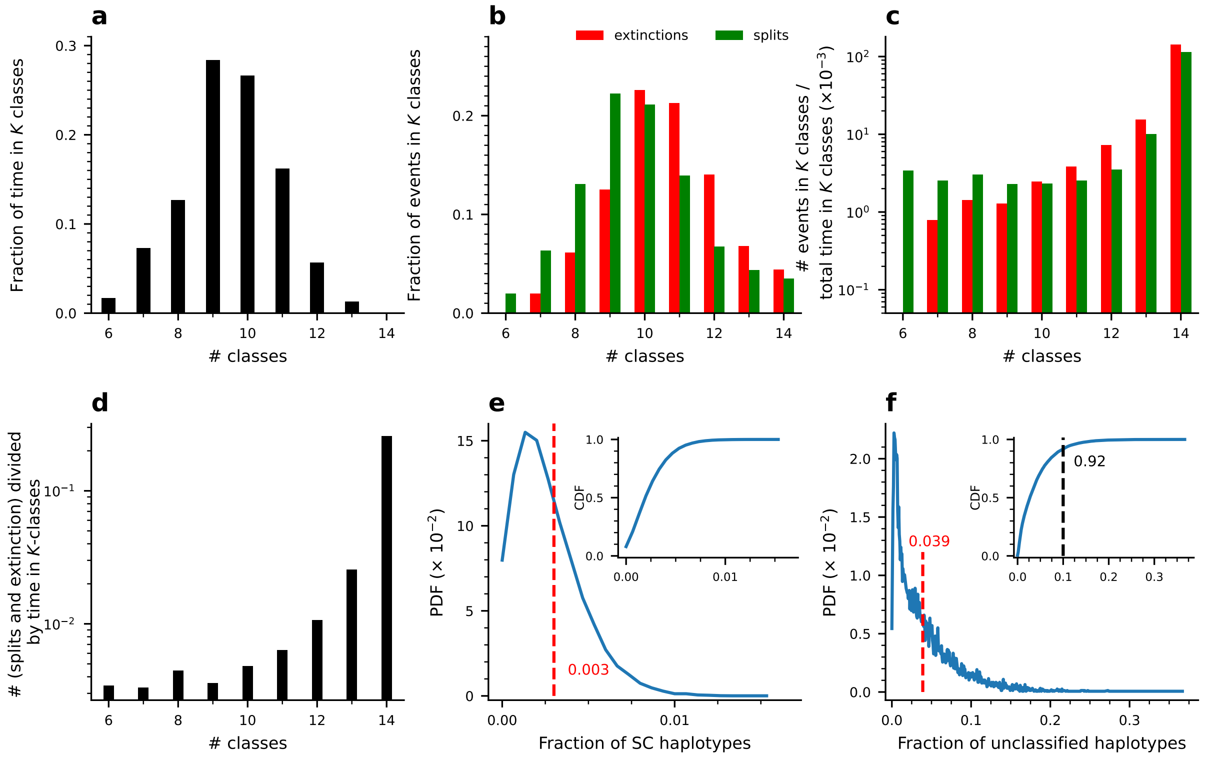


Figure S7: **The equilibrium number of classes increases when the population size increases, but the model retains its qualitative behavior, simulation results for $E_{\text{th}} = -6$, and $N = 750$.** **(a)** The fraction of time spent under $K$-classes state. **(b)** The fraction of split (green) and extinction (red) events that occurred under $K$-class state. **(c)** The extinction (split) rate – the number of extinction (red) and split (green) events in $K$-classes state divided by the time spent at this state – against $K$. (a-c) demonstrate a stable equilibrium, similar to Fig. 5, but with a higher equilibrium value of $K = 9$. Comparison to Fig. 5 where $N = 500$ was used, shows that a higher $N$ results in a higher number of classes, for the same energy threshold, $E_{\text{th}} = -6$. **(d)** Total event rate: sum of both split and extinction events under $K$ classes, divided by the time spent under $K$ classes (sum of the red and green bars as they appear in (c)). **(e)** Distribution of the proportion of self-compatible haplotypes, and **(f)** Distribution of the proportion of unclassified haplotypes across multiple simulation instances. The red dashed vertical lines in both panels show the distribution mean. The insets in both panels show the cumulative distributions. Results based on 13 independent runs, with a total of 156840 generations.

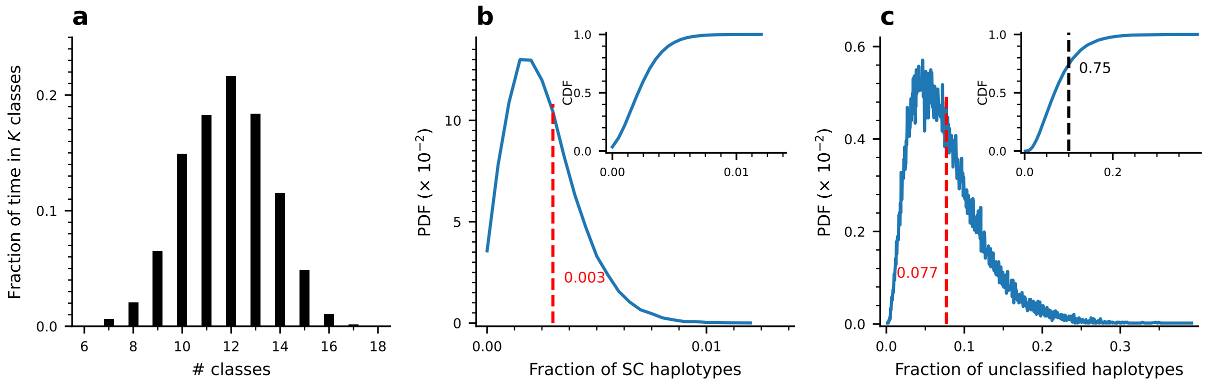


Figure S8: **The equilibrium number of classes increases when the population size increases, but the model retains its qualitative behavior, simulation results for $E_{\mathrm{th}} = -6$, and $N = 1000$.** **(a)** The fraction of time spent under $K$-classes state. The results demonstrate a stable equilibrium, similar to Fig. 5, but with a higher equilibrium value of $K = 12$ ,whereas for $N = 500$ the equilibrium class number was $K = 8$ (compare to Fig. 5). **(b)** Distribution of the fraction of self-compatible haplotypes. **(b)** Distribution of the fraction of unclassified haplotypes. Parameter values: $N = 1000$, $E_{\mathrm{th}} = -6$. Results based on 18 independent runs, with a total of 529400 generations

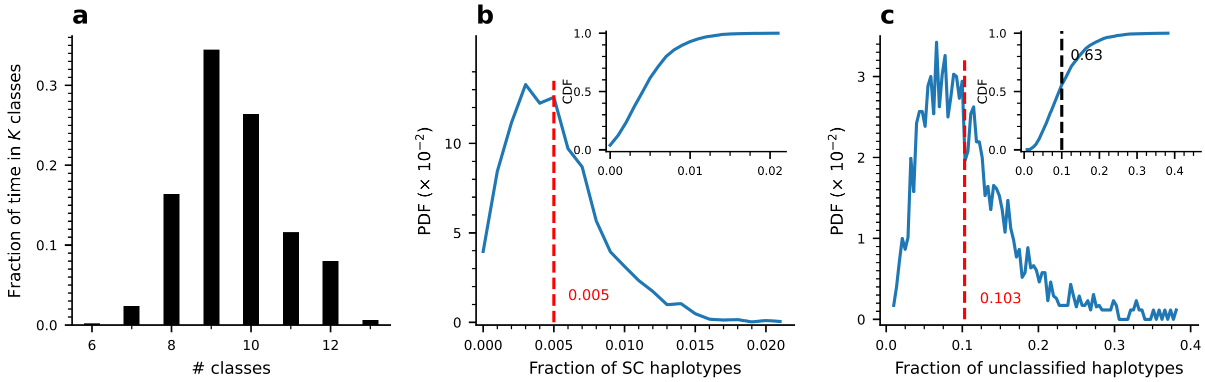


Figure S9: **The equilibrium number of classes, and the proportions of self-compatible and unclassified haplotypes increase with the mutation rate: simulation results with a higher mutation rate $p_{\mathrm{mut}} = 2 \times 10^{-4}$.** **(a)** The fraction of time spent in each $K$-class population state. The equilibrium class number here is $K = 9$ (compare to Fig. 5). **(b)** The proportion of self-compatible, and **(c)** the proportion of unclassified haplotypes are both higher under these parameters. Compare to Fig. S3 with a lower mutation rate of $p_{\mathrm{mut}} = 10^{-4}$. Results based on 9 independent runs, with a total of 40060 generations.

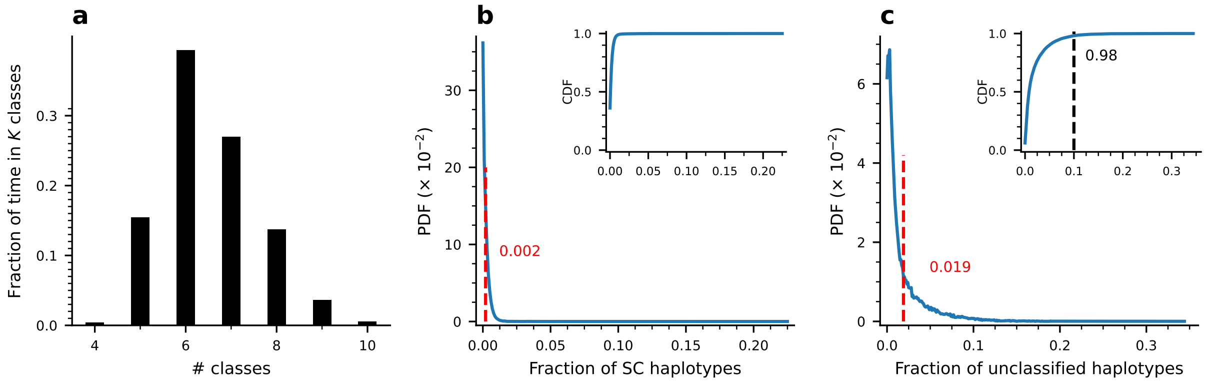


Figure S10: **The equilibrium number of classes, and the proportions of self-compatible and unclassified haplotypes increase with the mutation rate: simulation results with a lower mutation rate $p_{\text{mut}} = 0.5 \times 10^{-4}$.** **(a)** The fraction of time spent in $K$-class population state. The equilibrium class number here is $K = 6$ (compare to Fig. 5).**(b)** The proportion of self-compatible, and **(c)** the proportion of unclassified haplotypes are both lower under these parameters. Compare to Fig. S3 with a higher mutation rate of $p_{\text{mut}} = 10^{-4}$. Results based on 10 independent runs, with a total of 381710 generations.

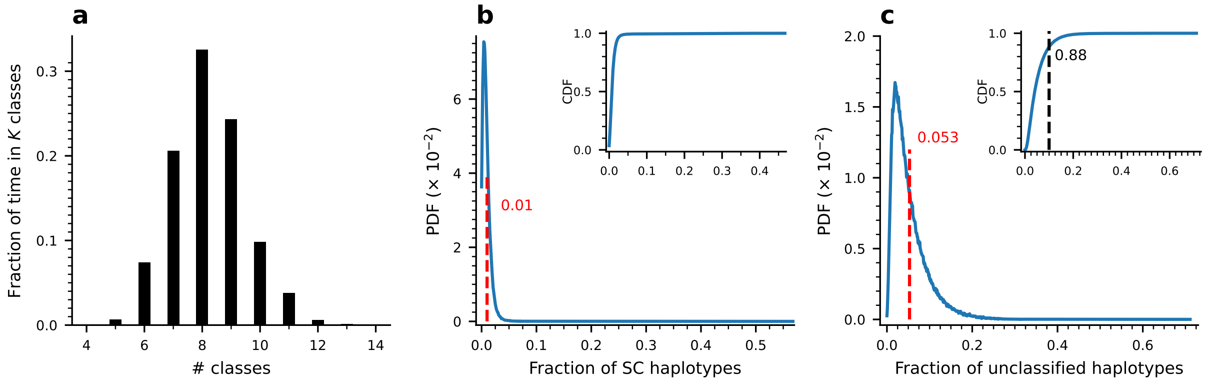


Figure S11: **The population proportions of self-compatible and unclassified haplotypes increase when inbreeding depression is relieved, but the equilibrium class number is not affected: simulation results for $\alpha = 0.8, \delta = 0.9$.** **(a)** The fraction of time spent in each $K$-class population state does not change (compare to 5). The population proportions of self-compatible **(b)**, and unclassified haplotypes **(c)** are higher under these parameters. Compare to Fig. S3 with $\delta = 1$ and $\alpha = 0.95$. Results based on 41 independent runs, with a total of 2102210 generations.

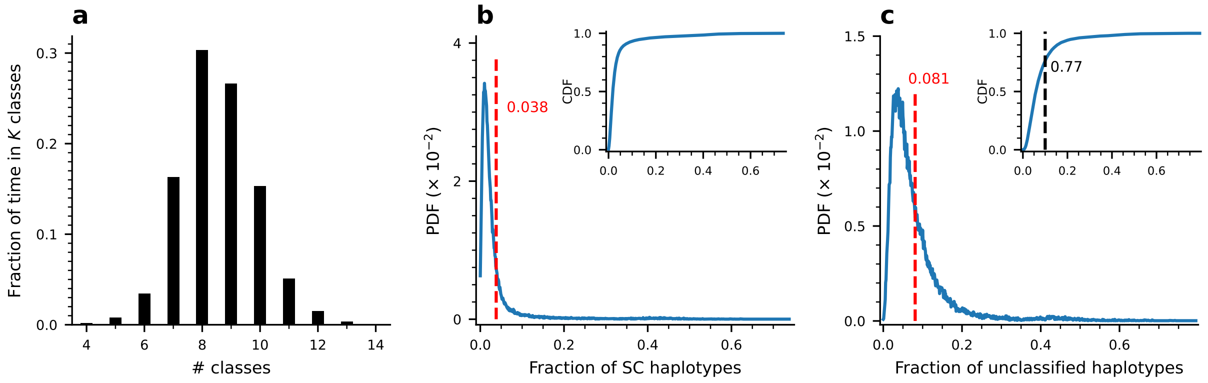


Figure S12: **The population proportions of self-compatible and unclassified haplotypes increase when inbreeding depression is relieved, but the equilibrium class number is not affected: simulation results for $\alpha = 0.6, \delta = 0.9$.** **(a)** The fraction of time spent in each $K$-class population state does not change (compare to 5). The population proportions of self-compatible **(b)**, and unclassified haplotypes **(c)** are higher under these parameters. Compare to Fig. S3 with $\delta = 1$ and $\alpha = 0.95$. Results based on 19 independent runs, with a total of 727740 generations.

.

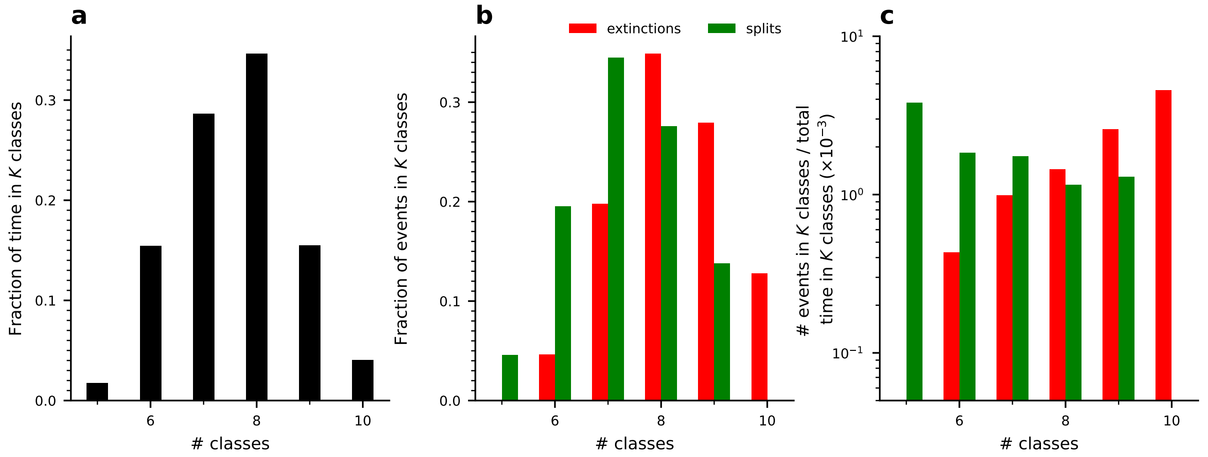


Figure S13: **Gene duplication and deletion have a negligible effect on the population structure and dynamics: dependence of split and extinction events on the present number of classes while gene deletion and duplication modules were disabled.** **(a)** The fraction of time spent under each $K$-class population state. **(b)** The fraction of split (green) and extinction (red) events that occurred under each $K$-class population state. **(c)** Extinction and split rates: the number of extinction (red) and split (green) events that occurred under each $K$-class state divided by the time spent at this state. These results are highly similar to those shown in Fig. 5 with $p_{\text{del}} = p_{\text{dup}} = 10^{-6}$. In particular, they suggest a stable equilibrium at an intermediate $K$ value, as before. Simulation results with $N = 500$, $E_{\text{th}} = -6$, $p_{\text{del}} = 0$, $p_{\text{dup}} = 0$. Results based on 11 independent runs, with a total of 704140 generations.

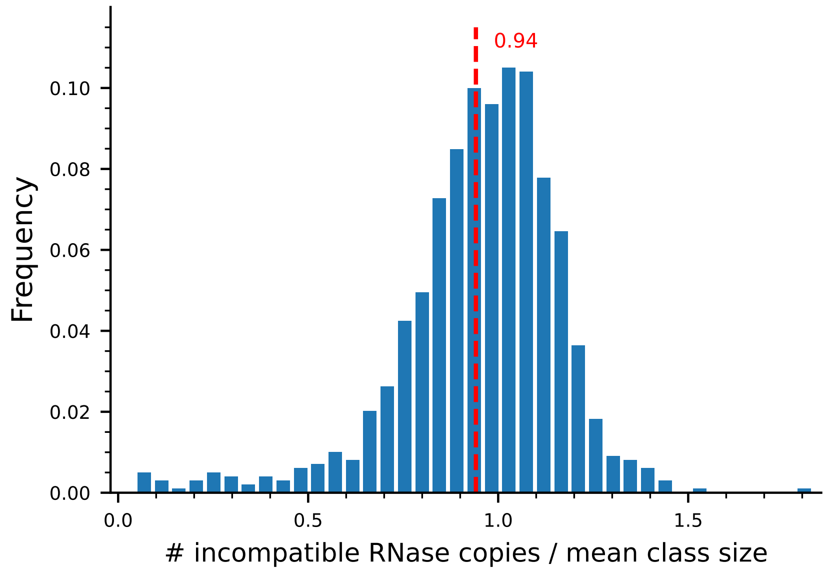


Figure S14: **Extinctions require that an incompatible RNase reaches a sizable copy number, but not necessarily an entire class – simulation results.** Here we show the number of extinction-driving incompatible RNases divided by the equilibrium class size at the moment of class extinction. Equilibrium class size was defined as $2N/K$, where $K$ is the number of population classes at that time. Dashed vertical line denotes the mean.

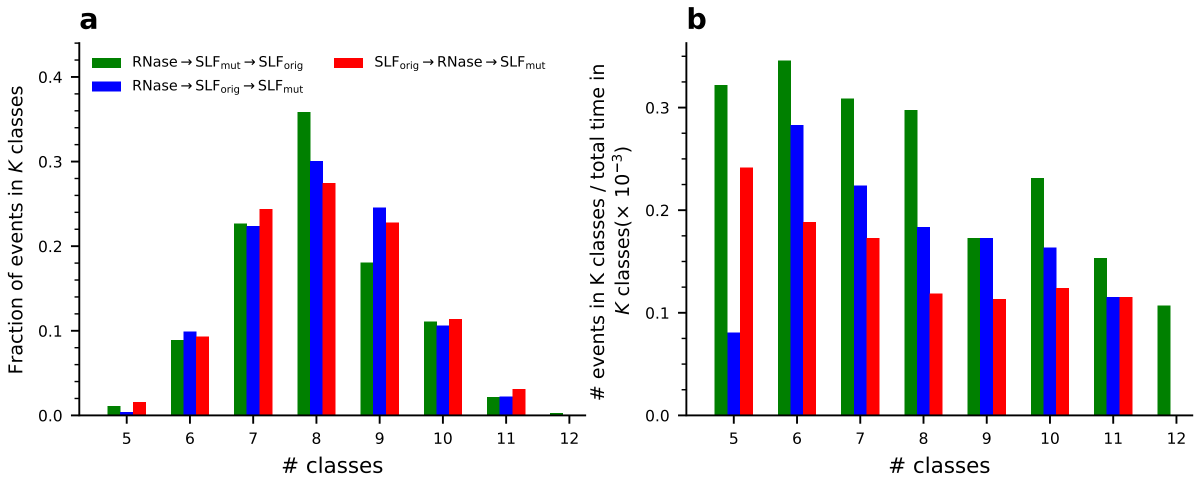


Figure S15: **The proportion and rate of split events against the number of population classes $K$, dissected by trajectory type.** **(a)** The proportion of events under each $K$-class population state for each of the three split trajectories, shown by different colors. **(b)** The rate of split events against the number of classes $K$ for each of the three trajectories. The event rate was calculated as the number of events divided by the time spent in that $K$-class population state.

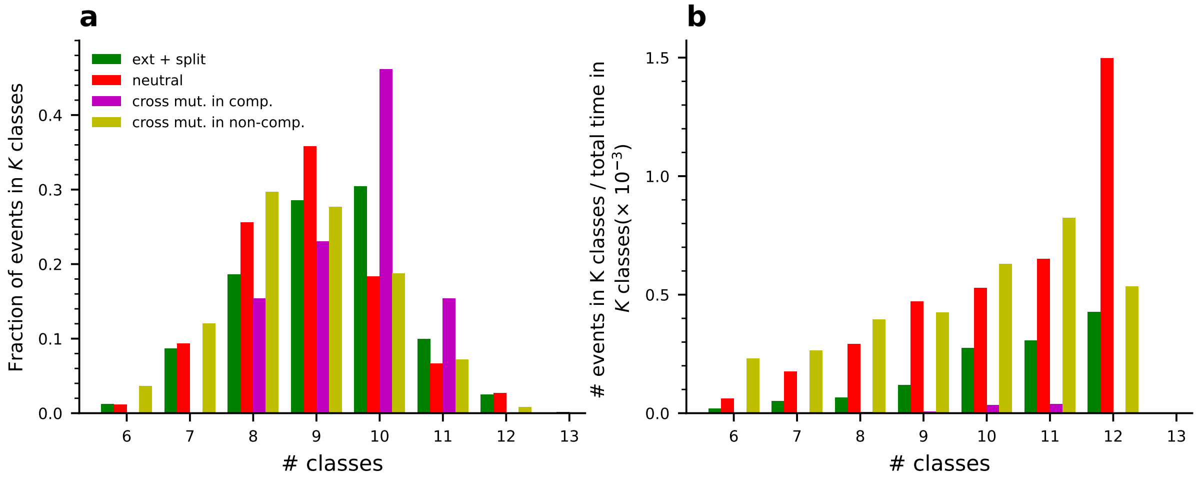


Figure S16: **The proportion and rate of extinction events against the number of population classes $K$, dissected by trajectory type.** **(a)** The proportion of events under each $K$-class population state for each of the different extinction trajectories, shown by different colors. **(b)** The rate of extinction events against the number of classes $K$ for each of the trajectories. The event rate was calculated as the number of extinction events divided by the time spent in that $K$-class population state. We observe that, for all trajectories the event rate increases with $K$.

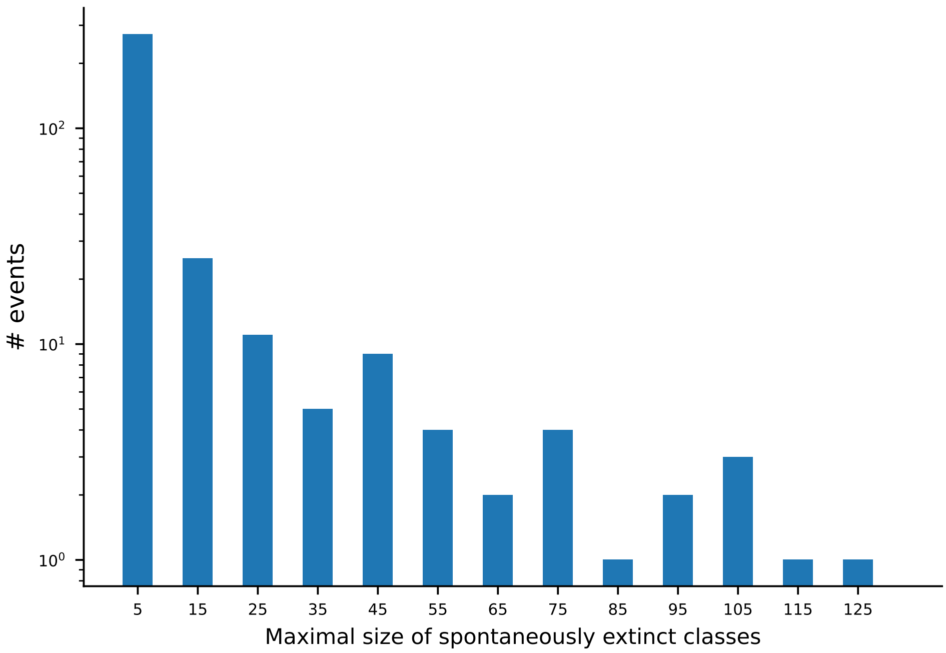


Figure S17: **The maximal size reached by classes that were later spontaneously extinct.** Spontaneous class extinctions occur with no involvement of any RNase mutation and hence are solely due to drift. Here we present a distribution of the maximal size reached by classes that later became spontaneously extinct. Clearly, the vast majority of these classes are very small (the distribution mean is approx. 10) and much smaller than the typical class size, which amounts to $>100$ for the equilibrium value of 8 to 9 classes. Since these classes could have occupied their maximal size long before their extinction, this could explain the few cases that exhibit a large maximal size.

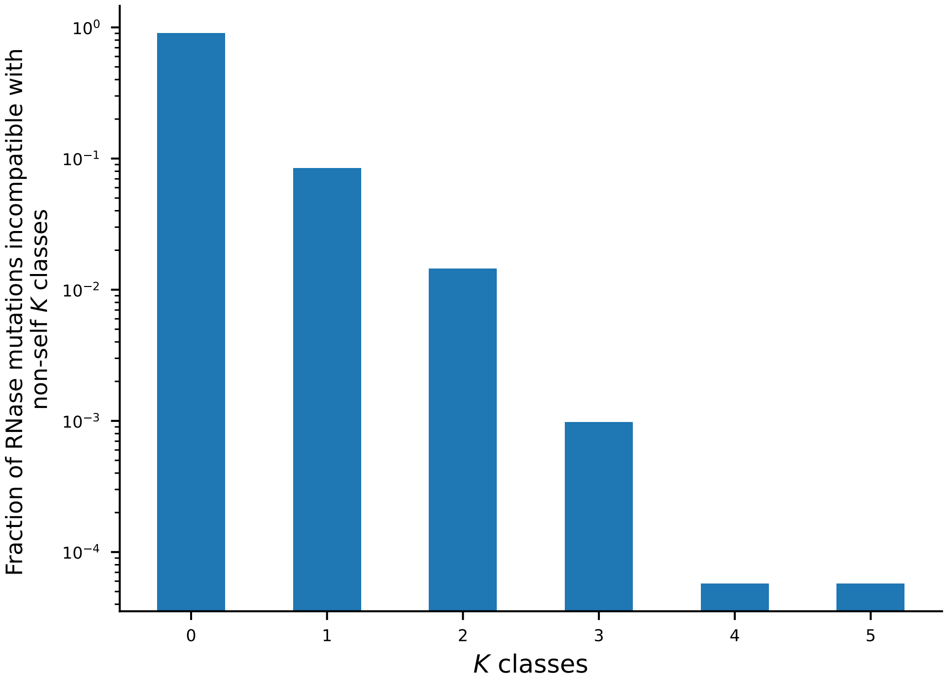


Figure S18: **The fraction of RNase mutations incompatible with non-self $K$ classes.** We tested the compatibility of all RNase mutations that survived more than 100 generations, regardless of whether they later caused a split, an extinction or none of these. We found that the vast majority of these mutations (90%) are compatible with all existing classes. Simulation parameters: $E_{\rm th} = -6$, $N = 500$. Exact values shown in the graph: [0.9003, 0.0841, 0.0145, 0.0010, 0.0001, 0.0001].
